# Loss of STIM2 in colorectal cancer drives growth and metastasis through metabolic reprogramming and PERK-ATF4 endoplasmic reticulum stress pathway

**DOI:** 10.1101/2023.10.02.560521

**Authors:** Trayambak Pathak, J. Cory Benson, Martin T. Johnson, Ping Xin, Ahmed Emam Abdelnaby, Vonn Walter, Walter A. Koltun, Gregory S. Yochum, Nadine Hempel, Mohamed Trebak

## Abstract

The endoplasmic reticulum (ER) stores large amounts of calcium (Ca^2+^), and the controlled release of ER Ca^2+^ regulates a myriad of cellular functions. Although altered ER Ca^2+^ homeostasis is known to induce ER stress, the mechanisms by which ER Ca^2+^ imbalance activate ER stress pathways are poorly understood. Stromal-interacting molecules STIM1 and STIM2 are two structurally homologous ER-resident Ca^2+^ sensors that synergistically regulate Ca^2+^ influx into the cytosol through Orai Ca^2+^ channels for subsequent signaling to transcription and ER Ca^2+^ refilling. Here, we demonstrate that reduced STIM2, but not STIM1, in colorectal cancer (CRC) is associated with poor patient prognosis. Loss of STIM2 causes SERCA2-dependent increase in ER Ca^2+^, increased protein translation and transcriptional and metabolic rewiring supporting increased tumor size, invasion, and metastasis. Mechanistically, STIM2 loss activates cMyc and the PERK/ATF4 branch of ER stress in an Orai-independent manner. Therefore, STIM2 and PERK/ATF4 could be exploited for prognosis or in targeted therapies to inhibit CRC tumor growth and metastasis.

**Highlights:**

- STIM2 regulates ER Ca^2+^ homeostasis independently of Orai and SOCE.
- STIM2 downregulation in colorectal cancer cells causes enhanced ER Ca^2+^ and is associated with poor patient prognosis.
- STIM2 downregulation induces PERK/ATF4 dependent ER stress in colorectal cancer.
- Increased ER stress drives colorectal cancer metabolic reprogramming, growth, and metastasis.

## Introduction

Colorectal cancer (CRC) is one of the leading causes of death in the United States, with an estimated 53,000 deaths per year (Siegel *et al*, 2020). A recent report showed an alarming annual increase in CRC incidence by 2.2 % in individuals younger than 50 years (Siegel *et al*, 2023). Despite recent advancements in detection and treatment, patients diagnosed with metastatic CRC have poor prognoses. Recent studies identified five CRC intrinsic subtypes (CRIS) based on their molecular and transcriptional signatures (Dunne *et al*, 2017; Guinney *et al*, 2015; Isella *et al*, 2017). Transcriptional reprogramming is required for cancer cells to overcome and adapt to environmental and physiological stressors and to evade death and undergo clonal selection and metastasis (Blancafort *et al*, 2005; Teng *et al*, 2020).

In healthy cells, altered function of the endoplasmic reticulum (ER) through exposure to stressors generally leads to increased accumulation of misfolded or unfolded proteins (Hetz, 2012). The increased unfolded protein burden in the ER triggers an adaptive response termed the unfolded protein response (UPR), which includes several homeostatic signaling pathways aimed at reducing unfolded proteins, recovering proper ER function, and sustaining cell survival. When cells fail to adapt during conditions of chronic ER stress, apoptotic cell death is triggered (Hetz, 2012). In cancer cells however, chronic UPR supports cancer cell adaptation and survival in response to various stress responses, including proteostasis defects, nutrient deprivation, and hypoxia. Once activated, the UPR promotes the generation of multiple transcription factors to support the transcriptional reprogramming of cancer cells (Corazzari *et al*, 2017; Costa-Mattioli & Walter, 2020; Hart *et al*, 2012; Tameire *et al*, 2019). There are three known branches of the UPR with unique signal transduction mechanisms that function in parallel to one another. Each branch is divided based on the presence of a specific endoplasmic reticulum (ER)-resident transmembrane protein: i) IRE1α (inositol requiring enzyme 1α), ii) PERK (double-stranded RNA-activated protein kinase (PKR)–like ER kinase), and iii) ATF6 (activating transcription factor 6) (Hetz, 2012; Walter & Ron, 2011). These three ER proteins bind the ER chaperone BiP from the ER luminal side, keeping them inactive under physiological conditions. Altered ER Ca^2+^ homeostasis has a direct effect on the ATP-ase cycle of the ER chaperone BiP and on BiP-substrate interactions, subsequently triggering the proteostatic effects of UPR (Preissler *et al*, 2020). Furthermore, activation of oncogenes such as cMyc also triggers bioenergetic processes such as protein synthesis and metabolic rewiring to support cancer cell growth and proliferation (Destefanis *et al*, 2020; Hart *et al*., 2012; Li *et al*, 2005; Morrish & Hockenbery, 2014; Pakos-Zebrucka *et al*, 2016; Satoh *et al*, 2017). Multiple studies showed that cMyc-induced transcriptional reprogramming activates the PERK/eIF2α/ATF4 branch of the UPR (Hart *et al*., 2012; Tameire *et al*., 2019).

Cytosolic, ER and mitochondrial Ca^2+^ are interconnected and are fueled and regulated through Ca^2+^ influx across the plasma membrane (PM) (Ben-Kasus Nissim *et al*, 2017; Emrich *et al*, 2021; Pathak *et al*, 2020; Pathak & Trebak, 2018; Trebak & Putney, 2017). In non-excitable cells, the major route of regulated Ca^2+^ influx into cells is the store-operated calcium entry (SOCE) mediated by PM-resident Orai channels (Emrich *et al*, 2022; Prakriya & Lewis, 2015; Trebak & Kinet, 2019; Trebak & Putney, 2017; Yoast *et al*, 2020a). SOCE is regulated by ER-resident Ca^2+^ sensing stromal-interacting molecules (STIM1 and STIM2), which are encoded by two distinct genes.

Stimulation of G protein-coupled receptors (GPCR) or receptor tyrosine kinase (RTK) that couple to isoforms of phospholipase C (PLC) leads to production of inositol-1,4,5-trsiphosphate (IP_3_). IP_3_ binds to IP_3_ receptor channels in the surface of the ER to cause Ca^2+^ release from the ER and subsequent ER Ca^2+^ store depletion. Upon ER Ca^2+^ depletion, STIM proteins aggregate and move into ER-PM junctions where they physically trap and activate Orai Ca^2+^ channels to trigger Ca^2+^ influx into the cytosol (Emrich *et al*., 2021; Yoast *et al*, 2020c). The N-terminal Ca^2+^-binding EF-hands of STIM1 and STIM2 have different affinities for Ca^2+^; STIM2 Kd for Ca^2+^ is ∼400 µM while STIM1 Kd for Ca^2+^ is ∼200 µM. As such, STIM2 activation is triggered by subtle depletion of ER Ca^2+^ stores while STIM1 requires substantial store depletion to be activated (Brandman *et al*, 2007b; Trebak & Kinet, 2019). Further, the C-terminal domains of STIM1 and STIM2 that interact with Orai are also distinct with STIM1 being a strong activator of SOCE while STIM2 is a weak activator (Emrich *et al*, 2019). While both STIM1 and STIM2 are concomitantly involved in SOCE across the range of agonist concentrations (Emrich *et al*., 2021), STIM2 function is mostly prominent under resting conditions and under moderate levels of receptor stimulation while STIM1 is the primary driver of receptor-activated SOCE during robust receptor activation (Brandman *et al*., 2007b; Emrich *et al*., 2021; Liou *et al*, 2005; Roos *et al*, 2005; Wedel *et al*, 2007). In summary, STIM2 and STIM1 are *bona fide* ER Ca^2+^ sensors and activators of SOCE, but they differ in their strength of activation of SOCE and operate at different ranges of ER Ca^2+^ store depletion and agonist concentrations.

Through the course of eukaryotic evolution, STIM appeared long after Orai, suggesting that STIM proteins likely functioned independently of Orai channels and only later did they co-opt an existing Orai for regulating SOCE (Collins & Meyer, 2011). Indeed, in addition to the established synergistic function of STIM1 and STIM2 proteins in regulating SOCE, several studies have reported on additional functions of STIM1 that are independent of SOCE. STIM1 was shown to regulate endothelial permeability by coupling to RhoA activation in a SOCE- and Orai1-independent manner(Shinde *et al*, 2013). In cardiomyocytes, STIM1 binds to phospholamban in the ER to indirectly activate SERCA2 and enhance sarcoplasmic reticulum (SR) Ca^2+^ levels (Zhao *et al*, 2015). STIM1 also supports the SR-PM functional coupling in contractile smooth muscle of Ca^2+^ release channels in the SR and ion channels on the PM (Krishnan *et al*, 2022). Furthermore, STIM1 was reported to regulate adenylyl cyclases (Motiani *et al*, 2018) and a number of PM ion channels and transporters, including transient receptor potential canonical (TRPC) channels, arachidonate-regulated Ca^2+^ (ARC) channels and PM Ca^2+^ ATPase (PMCA) (Gonzalez-Cobos *et al*, 2013; Ritchie *et al*, 2012; Shuttleworth *et al*, 2007; Worley *et al*, 2007; Zhang *et al*, 2013). However, SOCE-independent functions of STIM2 remain elusive, and the molecular mechanisms of STIM2-mediated activation of ER specific signaling pathways in either normal or cancer cells is unknown.

Here, we discovered that the loss of STIM2 expression is associated with poor colorectal cancer patient survival. we show a novel SOCE- and Orai-independent function of STIM2 in regulating ER Ca^2+^ homeostasis and UPR-dependent ER stress response in colorectal cancer. We demonstrate that loss of STIM2 in CRC alters ER Ca^2+^ levels and activates cMyc and the PERK/ATF4 branch of ER stress pathway to cause transcriptional reprogramming and metabolic rewiring of colorectal cancer cells. These changes in transcription and metabolism support the growth and metastasis of CRC cells in animal xenograft models. Therefore, we have discovered a previously unknown SOCE-independent function of STIM2 as a molecular switch regulating ER stress to control CRC survival and progression suggesting that STIM2 could be a therapeutic target of ER stress-induced CRC progression.

## Results

### Reduced STIM2 levels are associated with poor prognosis of colorectal cancer patients

We analyzed mRNA levels of STIM1 and STIM2 using publicly available data from The Cancer Genome Atlas (TCGA). STIM1 mRNA levels were significantly reduced, and STIM2 mRNA levels were unaltered in both colon and rectal adenocarcinoma (COAD-READ/COREAD) compared to unmatched normal tissue controls (**Figure 1A, B**). However, when we analyzed 20 colorectal tumor samples compared to their paired normal adjacent tissue obtained from patients at the Pennsylvania State University Hershey Medical Center, we observed a significant reduction in both STIM1 and STIM2 mRNA levels in the colorectal tumor samples compared to paired normal adjacent tissues (**Figure 1C**). Accessing TCGA data through UALCAN analysis portal (Chandrashekar *et al*, 2017) showed that Orai1, Orai2, and Orai3 mRNA levels were significantly upregulated in COAD patients (**Supplementary Figure 1I-K**). High STIM1 and low STIM2 expression were associated with TP53 mutation (**Supplementary Figure 1A, B**). However, we did not observe an association between STIM1 or STIM2 mRNA levels with mutations in phosphoinositide 3-kinases (PI3K), KRAS, or NRAS (**Supplementary Figure 1C-H**). Interestingly STIM2 mRNA, but not STIM1 mRNA levels correlated with high BRAF mutation (**Supplementary Figure 1G-H**). Further analysis revealed a significant reduction in STIM1 mRNA levels at all stages and slight upregulation of STIM2 at early (stage I/II) and no changes at later stages (stage III/IV) of COAD-READ patient samples (**Figure 1D, E**). Subsequently, Kaplan-Meier analysis showed a strong correlation between reduced STIM2 expression and poor prognosis in COAD or READ patients (**Figure 1F, G**). However, STIM1 expression did not correlate with the survival of COAD or READ patients (**Figure 1H, I**). Contrary to this, higher expression of Orai3 but not Orai1/2 was associated with poor prognosis in COAD patients (**Supplementary Figure 1L-N**). These data raise a very important question: how can two ER Ca^2+^ sensors, STIM1 and STIM2 that are established as synergistic activators of SOCE in healthy mammalian cell types, have differences in expression in colorectal cancer tissues and be associated with different outcomes of colorectal cancer patients? This suggests that the functions of STIM1 and STIM2 in colorectal cancer are quite unique, prompting us to explore the functions of STIM1 and STIM2 in colorectal cancer progression and whether their effects are mediated by SOCE.

**Figure 1.**
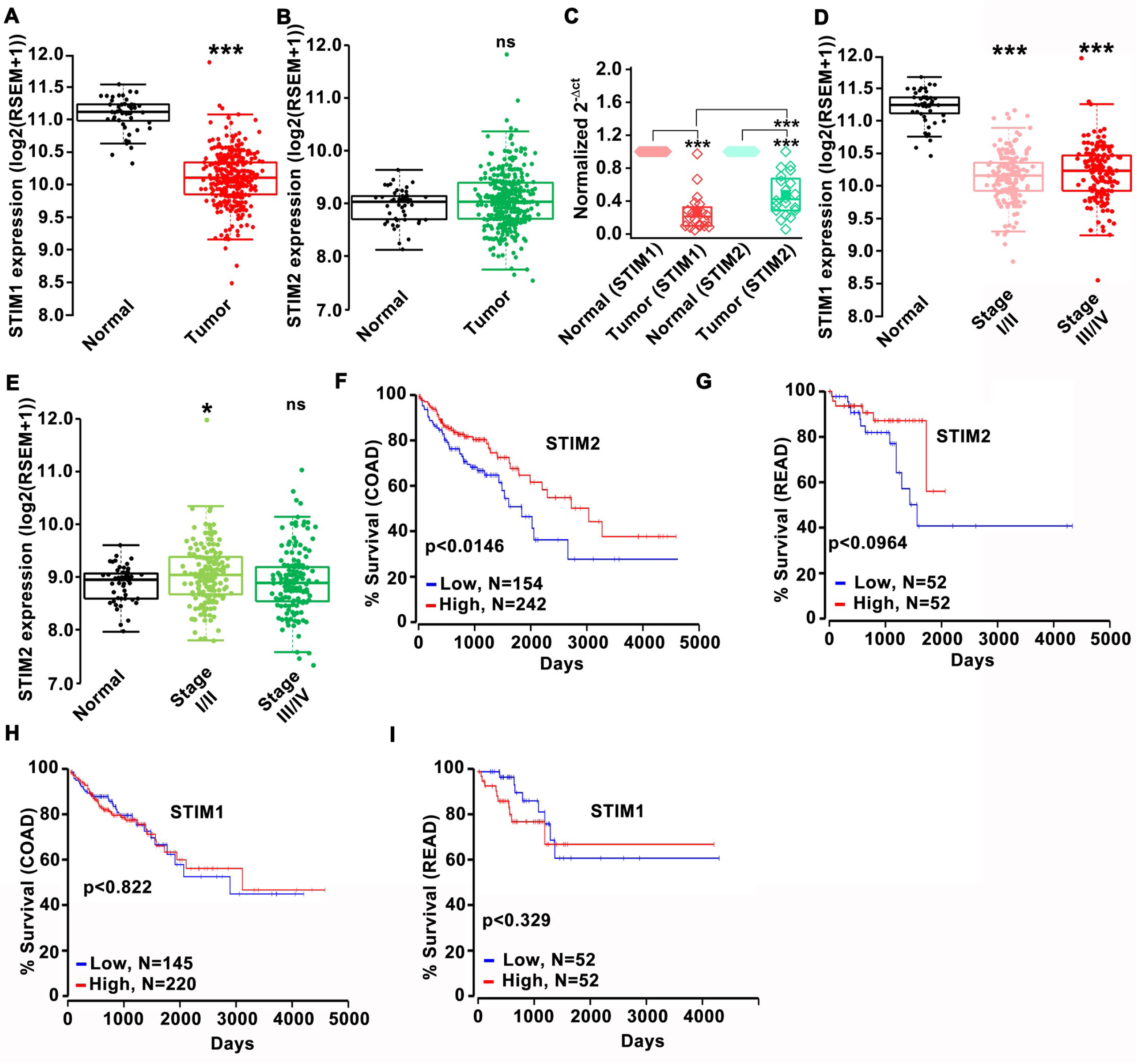
Low STIM2 expression in colorectal cancer is associated with poor patient prognosis. (A, B) TCGA data showing (A) STIM1 and (B) STIM2 mRNA levels in the unmatched tumor and adjacent normal tissue. (C) Analysis of STIM1 and STIM2 mRNA levels in matched tumor (n=20) and adjacent normal tissue (n=20) of CRC patients by RT-qPCR. The patient samples were obtained from Pennsylvania State University Hershey Medical Center. (D, E) TCGA data analysis of (D) STIM1 and (E) STIM2 mRNA levels in early (stage I/II) and late (III/IV) stages of CRC progression. (F, G) Kaplan plot of STIM2 showing survival of (F) colon adenocarcinoma (COAD) and (G) rectal adenocarcinoma (READ) patients. (H, I) Kaplan plot for STIM1 showing survival of (H) COAD and (I) READ patient. Kruskal-Wallis ANOVA was performed to test single variables between the two groups. *p<0.05, **p<0.01, and ***p<0.001.

### Loss of STIM2 promotes tumor growth and metastasis

Given that STIM1 and STIM2 are differentially expressed and that their expression levels show differences in association with CRC patient outcome, we investigated their functions in CRC. Thus, we focused our efforts on a comparative analysis of STIM1 and STIM2 in CRC progression and metastasis. We generated STIM1 knockout (STIM1 KO), STIM2 knockout (STIM2 KO), and STIM1/2 double KO (STIM DKO) cells from two human parental cell lines, HCT116 and DLD1 using CRISPR-Cas9 system (see methods for details). Sequence-specific guide RNAs for STIM1 and STIM2 were used to disrupt the gene and generate multiple knockout clones (see methods). Western blot analysis of wildtype HCT116, and their knockout clones: STIM1 KO #01, STIM1 KO #02, STIM2 KO #15, STIM2 KO #31, STIM DKO #22, and STIM DKO #34 showed complete loss of STIM1 and STIM2 proteins from their respective knockout clones (**Supplementary Figure 2A-C**). Similarly, western blot analysis of wildtype DLD1, and their knockout clones: STIM1 KO #03, STIM1 KO #05, STIM2 KO #04, STIM2 KO #09, STIM DKO #01, and STIM DKO #07 demonstrated complete loss of STIM1 and STIM2 proteins from their respective knockout clones (**Supplementary Figure 2D-F**). We did not observe any apparent compensation of STIM1 or STIM2 proteins in HCT116 STIM2 KO and STIM1 KO clones, respectively (**Supplementary Figure 2A-C**). However, a slight but significant reduction in STIM2 protein level was observed in DLD1 STIM1 KO clones (**Supplementary Figure 2D-F**). Fura-2 Ca^2+^ imaging was performed in each HCT116 and DLD1 knockout clone to functionally document STIM protein knockout. SOCE was measured in response to store depletion with 100 µM ATP in nominally Ca^2+^ free bath solution and subsequent addition of 2 mM Ca^2+^. HCT116 and DLD1 STIM1 KO and STIM DKO clones showed an almost complete abrogation of SOCE. HCT116 and DLD1 STIM2 KO clones exhibited a 50-70% reduction in peak SOCE compared to their respective controls, in line with our previous studies (Emrich *et al*., 2019), suggesting a significant contribution of STIM2 to SOCE in CRC cells (**Supplementary Figure 2G-J**)

Using these KO cells, we then utilized an intrasplenic xenograft model where luciferase-tagged parental HCT116, STIM1 KO #01, STIM2 KO #15, and STIM DKO #22 cells (5 × 10^5^ cells/mouse) were injected into the spleen of NOD-SCID mice (**Figure 2A**). Xenograft experiments were performed in two independent runs with 15 mice/group and mice were monitored weekly before being sacrificed at week 6. However, only the second run included mice injected with STIM DKO cells and therefore this group includes only 15 mice while the other three groups of mice (HCT116, STIM1 KO and STIM2 KO groups) include 30 mice/group. The *in vivo* metastasis and primary tumor growth were monitored through IP injection to mice with 100 µl luciferin and recording the total flux using In Vivo Imaging System (IVIS). We observed a significant increase in total luciferase luminescence flux in mice injected with HCT116 STIM2 KO #15 and STIM DKO #22 cells compared to parental HCT116 or STIM1 KO #01 cells (**Figure 2B, C**), suggesting that the loss of STIM2 enhances tumor growth. Furthermore, consistent with total flux measurement, the primary tumor weight at the time of sacrifice (6 weeks post-injection) was substantially higher in STIM2 KO #15 and STIM DKO #22 compared to HCT116 or STIM1KO #01 xenografted mice (**Figure 2D, E**).

**Figure 2.**
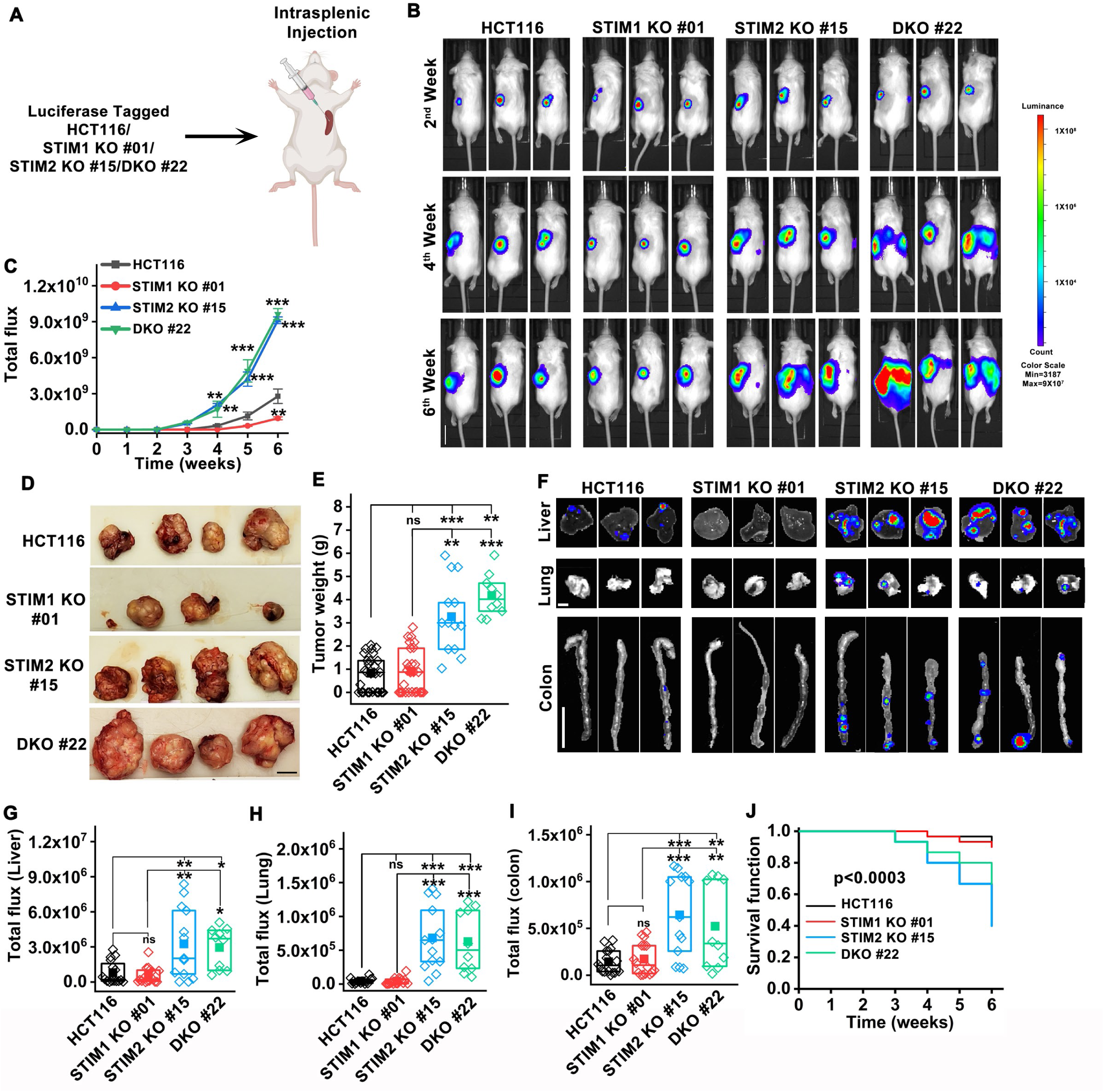
STIM2 loss enhances tumor growth and metastasis in mice. (A) Schematic representation of intrasplenic injection of luciferase tagged HCT116, STIM1 KO #01, STIM2 KO #15, and DKO #22 clones of HCT116 cells. The cells were injected at 5 X10^5^ cells/mice into the spleen of male NOD-SCID mice. (B, C) Male NOD-SCID mice injected with luciferase tagged HCT116, STIM1 KO #01, STIM2 KO #15, and DKO #22 clones of HCT116 cells (B) representative image with three mice per group, and (C) quantification of whole-body luminance (Scale bar 2.0 in). (D, E) Six weeks after intrasplenic injection of HCT116, STIM1 KO #01, STIM2 KO #15, and DKO #22 clones of HCT116 cells, the mice were sacrificed, and the primary tumor at the injection site was imaged, (D) representative image of primary tumors, and (E) quantification of primary tumor weight. (Scale bar 0.5 cm). (F-I) Post intrasplenic injection of HCT116, STIM1 KO #01, STIM2 KO #15, and DKO #22 clones of HCT116 cells, organs of intrasplenic injected mice were harvested, and luminance was measured (F) representative image of liver, lung, and colon showing metastasis, and quantification of total luminance in (G) liver (Scale bar 1 cm), (H) lung (Scale bar 0.5 cm), and (I) colon (Scale bar 0.5 cm). (J) Survival of NOD-SCID mice injected with HCT116, STIM1 KO #01, STIM2 KO #15, and STIM DKO #22 clones of HCT116 cells (p< 0.0003). ANOVA followed with a post hoc Tukey test to compare between groups. *p<0.05, **p<0.01 and ***p<0.001

A careful analysis of luciferase luminescence revealed that NOD-SCID mice xenografted with STIM2 KO #15 and STIM DKO #22 clones of HCT116 had luciferase activity other than in the primary tumor site, suggesting metastasis to distant organs (**Figure 2B**). Indeed, we observed a significantly higher luminescence in the liver, lung, and colon of STIM2 KO #15 and STIM DKO #22 compared to parental HCT116 or STIM1 KO #01 injected mice (**Figure 2F-I**). In mice injected with either parental HCT116 cells or STIM1 KO cells, out of the 28 (for HCT116) and 27 (for STIM1 KO) surviving mice at week 6, 16 mice/group showed metastasis to distant organs. For the case of mice injected with either STIM2 KO cells or STIM DKO cells, all surviving mice at week 6 (13 STIM2 KO-injected mice and 10 STIM DKO-injected mice) showed metastasis to distant organs. No significant difference was observed in luciferase luminescence in the kidney and heart of HCT116, STIM1 KO #01, STIM2 KO #15, and STIM DKO #22 cells (**Supplementary Figure 2K-M**). Survival analysis using the Kaplan Meier plot showed that NOD-SCID mice injected with STIM2 KO #15 and STIM DKO #22 cells had significantly reduced survival compared to HCT116 or STIM1 KO #01 injected mice (**Figure 2J**), suggesting that enhanced metastasis is the primary cause of lethality in STIM2 KO #15, and STIM DKO #22 xenografted mice. Collectively, these results show that loss of STIM2 enhances CRC tumor growth and metastasis.

### Deletion of STIM2 enhances spheroid formation, migration, and invasion of CRC cells

To delineate the mechanisms of increased tumor size and enhanced metastasis of STIM2 KO CRC cells, we performed spheroid formation, migration, and invasion assays. Parental cells, STIM1 KO, STIM2 KO, and STIM DKO clones of HCT116 and DLD1 cells were assessed for spheroid formation in ultra-low attachment round bottom 96-well plates. The cell-covered area was measured to determine the spheroid size. Within three days after spheroid formation, we observed a significant increase in spheroid size in STIM2 KO and STIM DKO clones compared to STIM1 KO clones and parental cells for both HCT116 and DLD1 (**Figure 3A-C**). On the other hand, we did not observe any significant difference in spheroid size between HCT116/DLD1 STIM1 KO clones and the parental cells (**Figure 3A-C**). Interestingly, we observed specifically in STIM2 KO and STIM DKO clones that some cells were loosely attached to the primary spheroid and were forming secondary spheroids (marked with red arrows, **Figure 3A**). Recent studies showed that cells that are loosely attached to the main spheroid body have higher migration and invasion potential (Stadler *et al*, 2018). Therefore, we investigated the migration and invasion properties of STIM1 KO, STIM2 KO, and STIM DKO clones of HCT116 and DLD1. We performed the gap closure assay and the Boyden chamber assay as means to quantify contact dependent migration and chemotaxis-dependent migration and invasion, respectively (Decaestecker et al., 2007; Guy et al., 2017). We found a significant increase in gap closure in STIM1 KO, STIM2 KO, and STIM DKO clones of both HCT116 and DLD1 cells at 24 hr compared to controls (**Supplementary Figure 3E-H**). Interestingly, when we performed migration and invasion assays using Boyden chambers (Matrigel-coated for invasion assays) we revealed a significant and selective increase in migration and invasion of STIM2 KO and STIM DKO clones of HCT116 and DLD1 cells compared to their respective controls (**Figure 3D-G**). However, the migration and invasion of STIM1 KO of HCT116 and DLD1 cells were either significantly reduced or unaffected compared to their respective parental cell controls (**Figure 3D, G**), suggesting a specific effect of STIM2 loss on chemotaxis-dependent migration and invasion. Consistent with increased invasion, GSEA analysis showed that hallmark genes of Endothelial Mesenchymal transition (EMT) were significantly enriched in STIM2 KO and STIM DKO cells (**Figure 5**), with matrix metalloproteases 16 (MMP16) transcript being one of the EMT genes significantly upregulated in all clones of HCT116 cells following STIM2 KO and STIM DKO. This was validated in STIM2 KO and STIM DKO of HCT116 and DLD1 cells compared to control cells (**Figure 3 H and J**). However, we observed a significant decrease in MMP14 mRNA levels in STIM1 KO, STIM2 KO, and STIM DKO of HCT116 and DLD1 cells compared to control cells (**Figure 3I and K**), indicating that MMP16 may be one of the regulators of invasion in STIM2 KO and STIM DKO colorectal cancer cells.

**Figure 3.**
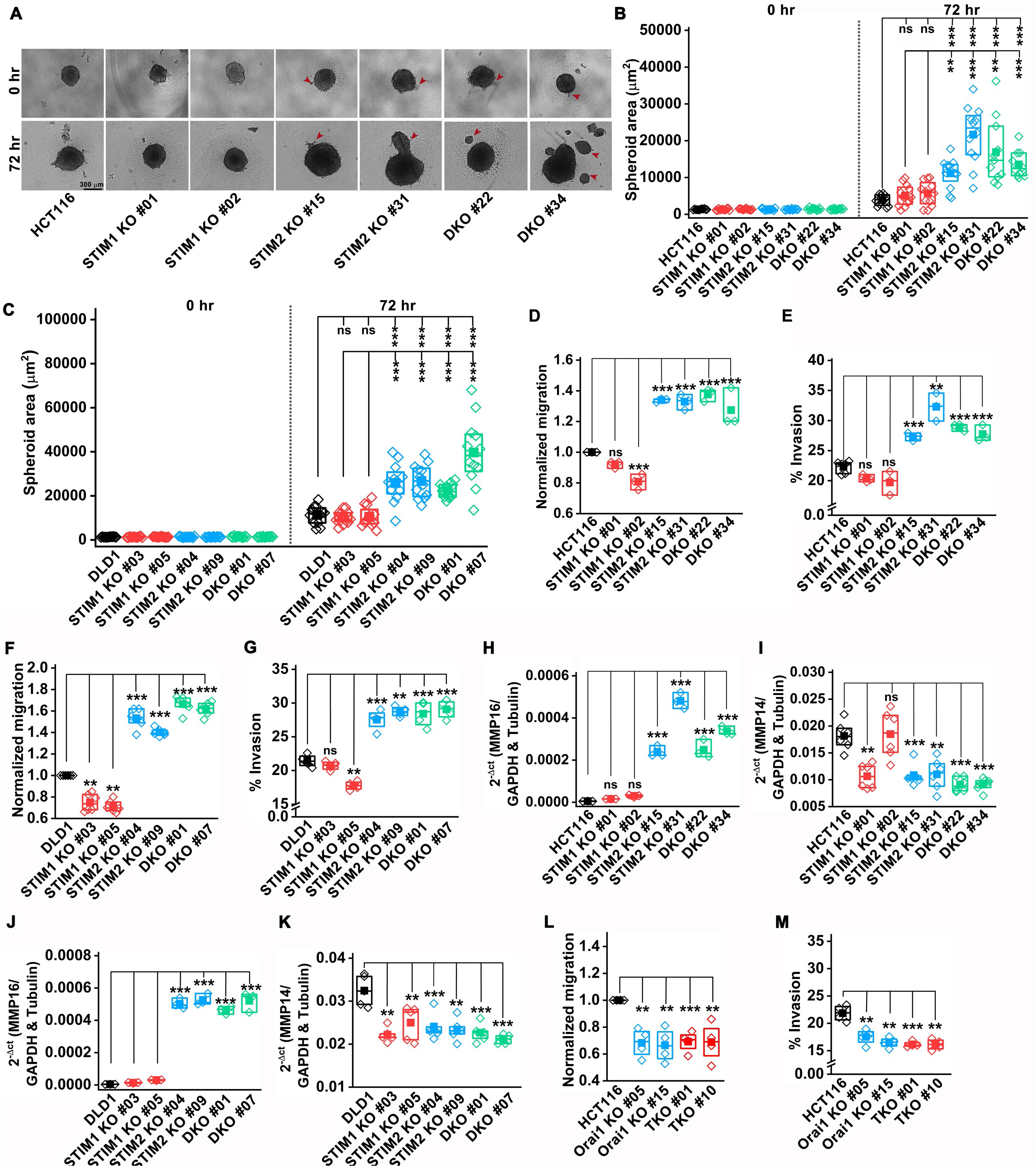
Loss of STIM2 enhances CRC proliferation and metastasis. (A, B) Spheroid formation assay showing (A) representative image of spheroid formed by HCT116, and clones of STIM1 KO, STIM2 KO, and DKO in HCT116 cells, (B) quantification of the spheroid area at 0 and 72 hr. (Scale bar 300 µm) (C) Quantification of the spheroid area of DLD1 and clones of STIM1 KO, STIM2 KO, and DKO in DLD1 cells. (D, E) Quantification of (D) normalized migration and (E) percent invasion of HCT116 and clones of STIM1 KO, STIM2 KO, and DKO in HCT116 cells using Boyden chamber assay. (F, G) Quantification of (F) normalized migration and (G) percent invasion of DLD1 and clones of STIM1 KO, STIM2 KO, and DKO in DLD1cells using Boyden chamber assay. (H, I) RT-qPCR analysis of mRNA levels of (H) MMP16 and (I) MMP14 in HCT116, STIM1 KO, STIM2 KO, and DKO clones of HCT116 cells. (J, K) RT-qPCR analysis of mRNA levels of (J) MMP16 and (K) MMP14 in DLD1 and clones of STIM1 KO, STIM2 KO, and DKO clones of DLD1 cells. (L, M) Quantification of (L) normalized migration and (M) percent invasion of HCT116 and clones of Orai1 KO, and Orai TKO in HCT116 cells using Boyden chamber assay. ANOVA followed with a post hoc Tukey test except for F-M where the paired t-test was used to compare between groups. *p<0.05, **p<0.01 and ***p<0.001

When we performed proliferation assays using the CyQUANT dye, we observed a substantial reduction in proliferation of all STIM1 KO, STIM2 KO, and STIM DKO clones of HCT116 and DLD1 cells compared to control at 48 hr and 72 hr in culture (**Supplementary Figure 3A, B**). Nevertheless, at 72 hr, the proliferation of STIM2 KO and STIM DKO clones was significantly higher than the STIM1 KO clones of HCT116 and DLD1 (**Supplementary Figure 3A, B**). We also performed *in vitro* colony formation assays to determine cell survival. Similar to proliferation assays, there was a significant decrease in the number of colonies formed by STIM1 KO, STIM2 KO, and STIM DKO clones of HCT116 cells compared to control cells. Nevertheless, the number of colonies formed by STIM2 KO and STIM DKO cells was significantly higher than that of STIM1 KO cells (**Supplementary Figure 3C, D**).

Given that the loss of STIM1 and STIM2 expression differentially affect COREAD cell metastatic behavior, suggests that their action is SOCE independent. To unequivocally rule out the involvement of SOCE in the increased CRC migration and invasion triggered by the loss of STIM2, we generated Orai1 KO clones and Orai1, Orai2, and Orai3 triple KO (Orai TKO) clones of HCT116 cells using the CRISPR-Cas9 system. RT-qPCR analysis showed the absence of Orai1 mRNA from Orai1 KO and the absence of Orai1, Orai2, and Orai3 mRNA from Orai TKO clones of HCT116 (**Supplementary Figure 3I-K**). As expected, Fura-2 Ca^2+^ imaging showed almost complete loss of SOCE in HCT116 Orai1 KO and Orai TKO cells (**Supplementary Figure 3L, M**). As compared to control cells, Orai1 KO and Orai TKO clones of HCT116 showed a significant increase in gap closure in wound healing assays in a manner reminiscent of all STIM knockout clones (**Supplementary Figure 3N-P**). However, unlike STIM2 KO and STIM DKO cells, Orai1 KO and Orai TKO cells showed decreased migration and invasion in the Boyden chamber assays (**Figure 3L, M**). These data strongly argue that the effect of STIM2 loss on chemotaxis-mediated CRC migration and invasion is Orai- and SOCE-independent.

### Loss of STIM2 leads to CRC mitochondrial remodeling and metabolic reprogramming

Previous studies in primary T cells, B cells, airway smooth muscle cells and astrocytes have shown that STIM1 is required for antigen- and growth factor-mediated induction of glycolysis and mitochondrial metabolism (Emrich *et al*, 2023; Johnson *et al*, 2022; Kaufmann *et al*; Novakovic *et al*, 2023; Vaeth *et al*, 2017). Therefore, we performed whole-cell metabolomics on parental HCT116, STIM1 KO #01, STIM2 KO #15, and STIM DKO #22 clones of HCT116 cells and observed a marked change in polar metabolites between STIM1 KO, STIM2 KO, and STIM DKO clones compared to control HCT116 cells (**Supplementary Figure 4A-C**). Interestingly, the STIM2 KO #15 and STIM DKO #22 clones showed a significant alteration in polar metabolites compared to the STIM1 KO #01 clone of HCT116 cells (**Supplementary Figure 4D, E**). Further analysis of metabolomics data showed that the metabolites of the glycolysis pathway were upregulated in all three replicates of STIM2 KO #15 and STIM DKO #22 clones compared to STIM1 KO #01 and HCT116 control (**Figure 4A**). Since perturbation of STIM2 function led to upregulation of glycolytic metabolites, we used Seahorse XFe24 to measure the extracellular acidification rate (ECAR) of all HCT116 and DLD1 cell variants. Consistent with the metabolomics data, the glycolytic capacity of STIM2 KO and STIM DKO clones was significantly increased. However, glycolysis was significantly decreased in STIM1 KO clones of HCT116 and DLD1 as well as in Orai1 KO and Orai TKO clones of HCT116 compared to their respective controls (**Figure 4B, and Supplementary Figure 5A, B**). Furthermore, we measured glucose and lactate in growth media using the YSI bioanalyzer system and observed a significant increase in glucose consumption and lactate generation by STIM2 KO and STIM DKO clones of HCT116 cells (**Figure 4C, D**). In contrast, STIM1 KO clones of HCT116 cells showed a significant decrease in glucose consumption and lactate generation compared to control (**Figure 4C, D**), suggesting that STIM2 regulates the metabolic transformation of CRC cells in a SOCE-independent manner. The RNA sequencing analysis from STIM1 KO, STIM2 KO, and STIM DKO clones of HCT116 revealed that Solute carrier family 2 A1 (SLC2A1), also known as glucose transporter member 1 (GLUT1), was significantly upregulated in STIM2 KO and STIM DKO clones compared to STIM1 KO or HCT116 (**Figure 5A-D**). Western blot analysis determined that, indeed, the protein levels of GLUT1 were significantly upregulated in STIM2 KO and STIM DKO clones of HCT116 and DLD1 cells (**Figure 4G, H, and Supplementary Figure 5C, D**). There was no change in GLUT1 protein levels in STIM1 KO clones of HCT116 and DLD1 cells and in Orai1 KO and Orai TKO clones of HCT116 cells compared to their respective controls (**Figure 4G, H, and Supplementary Figure 5E, F**), suggesting that increased GLUT1 levels supports the increased ECAR, glucose consumption, and lactate generation of STIM2 KO cells. These data indicate that STIM2 regulates glycolysis of colorectal cells independently of SOCE.

**Figure 4.**
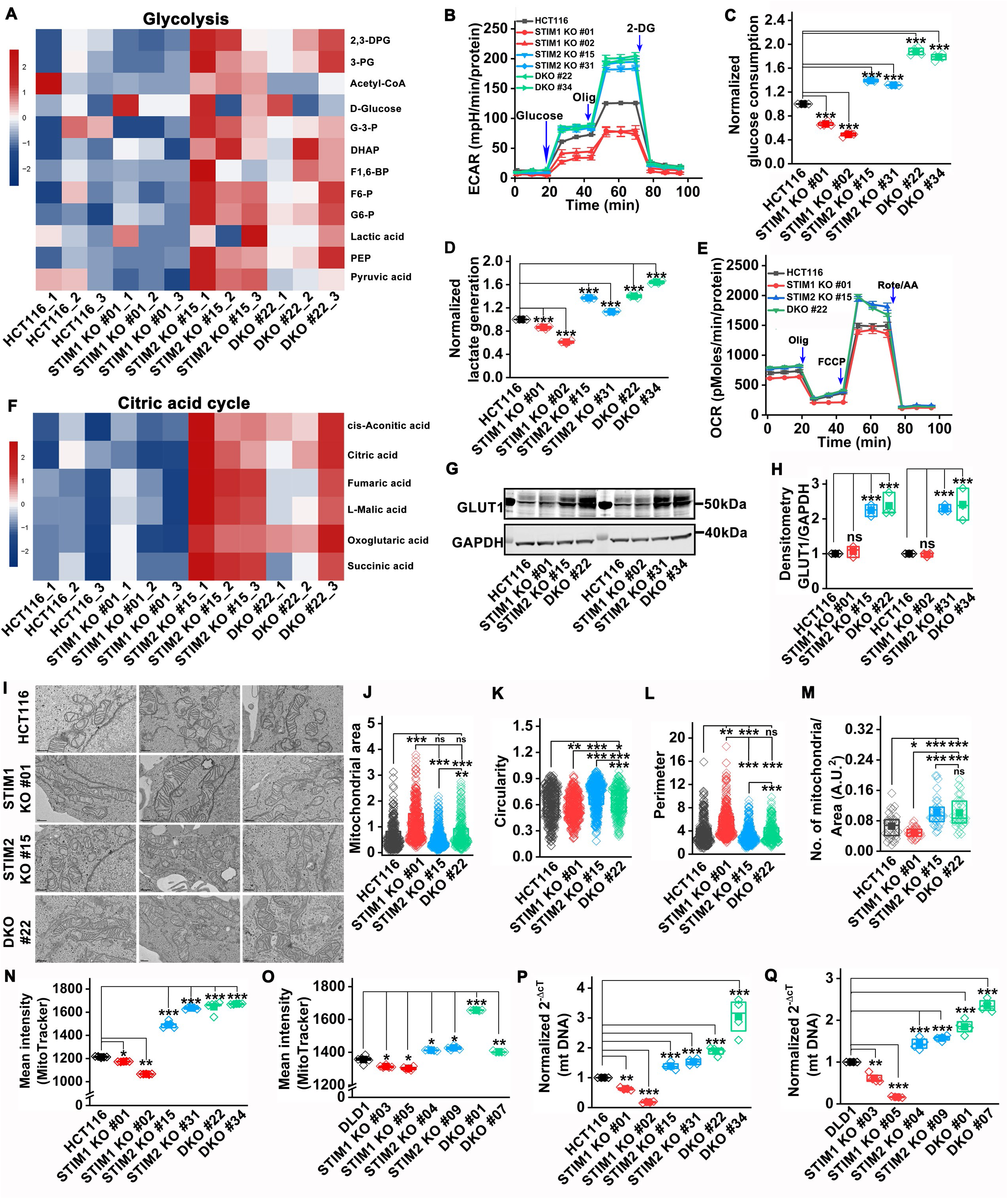
STIM2 regulates mitochondrial biogenesis and metabolism in CRC cells. (A) Heatmap showing significantly increased levels of glycolysis pathway metabolites in HCT116 STIM2 KO #15 and HCT116 DKO #22 compared to HCT116, and HCT116 STIM1 KO #01STIM1 KO #01 cells. (B) Extracellular acidification rate (ECAR) of HCT116 and clones of STIM1 KO, STIM2 KO, and DKO in HCT116 cells. (C, D) Measurement of (C) glucose consumption and (D) lactate generation in HCT116 and clones of STIM1 KO, STIM2 KO, and DKO in HCT116 cells. (E) Oxygen consumption rate (OCR) in HCT116 and STIM1 KO #01, STIM2 KO #15, and DKO #22 clones of HCT116 cells. (F) Heatmap showing significantly increased levels of citric acid cycle metabolites in HCT116 STIM2 KO #15 and HCT116 DKO #22 compared to HCT116, and HCT116 STIM1 KO #01 cells. (G, H) Measurement of GLUT1 protein level (G) representative western blot probed with anti-GLUT1 and GAPDH antibody and (H) densitometric analysis of GLUT1 normalized to GAPDH in HCT116 and clones of STIM1 KO, STIM2 KO, and DKO in HCT116 cells. (I-M) Ultrastructure of mitochondria analyzed by electron microscopy, (I) representative electron micrograph showing mitochondrial structure, measurement of (J) area, (K) circularity, (L) perimeter, and (M) mitochondrial number in HCT116 and STIM1 KO #01, STIM2 KO #15, and DKO #22 colons of HCT116 cells. (N, O) Flow cytometric analysis of MitoTracker intensity in (N) HCT116 and clones of HCT116 STIM1 KO, HCT116 STIM2 KO, and HCT116 DKO, and (O) DLD1, and clones of DLD1 STIM1 KO, DLD1 STIM2 KO, and DLD1 DKO cells. (P, Q) RT-qPCR analysis of mitochondrial DNA (mtDNA) in (P) HCT116 and clones of HCT116 STIM1 KO, HCT116 STIM2 KO, and HCT116 DKO, and (Q) DLD1, and clones of DLD1 STIM1 KO, DLD1 STIM2 KO, and DLD1 DKO cells. Statistical significance was calculated using ANOVA followed by a post hoc Tukey test to compare multiple groups except for C, D, H, P and Q, where the paired t-test was used to compare between groups. *p<0.05, **p<0.01 and ***p<0.001

The metabolic analysis also showed a significant increase in citric acid cycle metabolites in STIM2 KO #15 and STIM DKO #22 compared to STIM1 KO #01 clones and HCT116 cells (**Figure 4F**). Therefore, we determined the oxygen consumption rate (OCR) of all HCT116 and DLD1 cell variants using Seahorse assays. We observed a significant increase in maximal respiratory capacity in STIM2 KO and STIM DKO clones of HCT116 and DLD1 cells compared to control cells (**Figure 4E and Supplementary Figure 5G**). Conversely, the maximal respiratory capacity was significantly reduced in STIM1 KO clones of HCT116 and DLD1 cells and in Orai1 KO and Orai TKO clones of HCT116 cells (**Figure 4E and Supplementary Figure 5G, H**). Further, we assessed the protein levels of mitochondrial respiratory complex I-V to determine whether increased OCR in STIM2 KO and STIM DKO clones was due to increased protein levels of these complexes. Interestingly, we did not observe a significant change in protein levels of NADH dehydrogenase [ubiquinone] 1 β subcomplex subunit 8 (NDUFB8, Complex-I), Succinate dehydrogenase [ubiquinone] iron-sulfur subunit (SDHB, Complex-II), Cytochrome b-c1 complex subunit 2 (UQCRC2, Complex-III), Cytochrome c oxidase subunit 1 (MTCO1, Complex IV), and ATP synthase subunit α (ATP5A, Complex-V) in STIM1 KO, STIM2 KO and STIM DKO clones of HCT116 cells (**Supplementary Figure 5I-N**). We measured the mitochondrial and cytosolic superoxide levels by staining cells with MitoSOX or CellROX respectively. There was no significant change in mitochondrial and cytosolic superoxide levels in STIM1 KO, STIM2 KO and STIM DKO clones of HCT116 cells (**Supplementary Figure 5O, P**). Mitochondrial membrane potential measured by TMRE staining showed no difference in mitochondrial membrane potential between STIM1 KO, STIM2 KO and STIM DKO clones of HCT116 cells (**Supplementary Figure 5Q**). Multiple studies reported that the pentose phosphate pathway (PPP) is required for nucleotide biosynthesis, and plays a major role in metabolism, proliferation, and chemoresistance of colorectal cancer (Gao *et al*, 2019; Jiang *et al*, 2014; Lin *et al*, 2022; Patra & Hay, 2014). Our metabolomic analysis showed a significant increase in PPP metabolites in STIM2 KO #15, STIM DKO #22 clones, compared to STIM1 KO #01 or HCT116 control cells (**Supplementary Figure 5R**), suggesting that the loss of STIM2 enhances PPP to support colorectal cancer growth. Collectively these data show that loss of STIM2 positively regulates CRC cell metabolic activity.

Because changes in overall cell metabolism are typically associated with changes in either mitochondrial structure, number, or function, we performed transmission electron microscopy (TEM) imaging to assess any changes in mitochondrial number and structure. The single mitochondrion area and perimeter were slightly decreased, and mitochondria were slightly more circular in STIM2 KO #15 and STIM DKO #22 clones compared to STIM1 KO #01 clone of HCT116 (**Figure 4I-L)**. However, we found a significant increase in mitochondrial number in STIM2 KO #15, STIM DKO #22 clones, and a decrease in mitochondrial number in STIM1 KO #01 with longer single mitochondria compared to HCT116 control (**Figure 4I, M**). We stained cells with MitoTracker green to further examine the mitochondrial number with flow cytometry. We found a significant increase in MitoTracker intensity in STIM2 KO and STIM DKO clones of HCT116 and DLD1 cells compared to control cells (**Figure 4 N, O**). The MitoTracker intensity was significantly decreased in STIM1 KO clones of HCT116 and DLD1 compared to control cells (**Figure 4 N, O**). Mitochondrial DNA, as measured by qPCR was markedly increased in STIM2 KO and STIM DKO clones and significantly decreased in STIM1 KO clones of HCT116 and DLD1 cells compared to control (**Figure 4 P, Q**).

### Loss of STIM2 promotes transcriptional reprogramming of colorectal cancer cells

We performed transcriptional profiling of parental cells, STIM1 KO, STIM2 KO, and STIM DKO clones of HCT116 using RNA sequencing. Approximately 10,000 genes emerged as common hits, which were differentially expressed between HCT116 cell variants (**Figure 5A-D and Supplementary Figure 6A-D**). Gene set enrichment analysis (GSEA) revealed that the gene set involved in cMyc regulation, Oxidative phosphorylation (OXPHOS), and unfolded protein response (UPR) were upregulated in STIM2 KO and STIM DKO clones of HCT116 versus control. Among the Hallmark gene sets, cMyc target V1, V2, OXPHOS, Endothelial to mesenchymal (EMT), G2M, and E2F targets were positively regulated in STIM2 KO and STIM DKO clones of HCT116 (**Figure 5 E-N Supplementary Figure 7 A-C**). We aligned differentially expressed genes to the Kyoto Encyclopedia of Genes and Genomes (KEGG) pathway analysis and found that ribosome biogenesis, RNA transport, and translation pathways were significantly upregulated in STIM2 KO and STIM DKO clones of HCT116 (**Supplementary Figure 6E-J**), suggesting increased translation and protein accumulation. We did not observe an enrichment of translation pathways in STIM1 KO clones of HCT116 cells (**Supplementary Figure 6E, F**). We also performed gene ontology (GO) enrichment analysis to understand the distribution of differentially expressed genes in biological processes (GO-BP). Like the KEGG analysis, GO-BP analysis also showed significant enrichment of ribosome biogenesis, RNA metabolism, and translation initiation enrichment in STIM2 KO and STIM DKO clones of HCT116 as compared to HCT116 control (**Supplementary Figure 6K-P**). Collectively, these data show that deletion of STIM2 upregulates ribosome biogenesis and translation, suggesting enhanced UPR and increased protein accumulation. Importantly, the loss of STIM2 affects completely different pathways compared to that of STIM1, lending further support to a SOCE-independent function of STIM2 in CRC progression.

**Figure 5.**
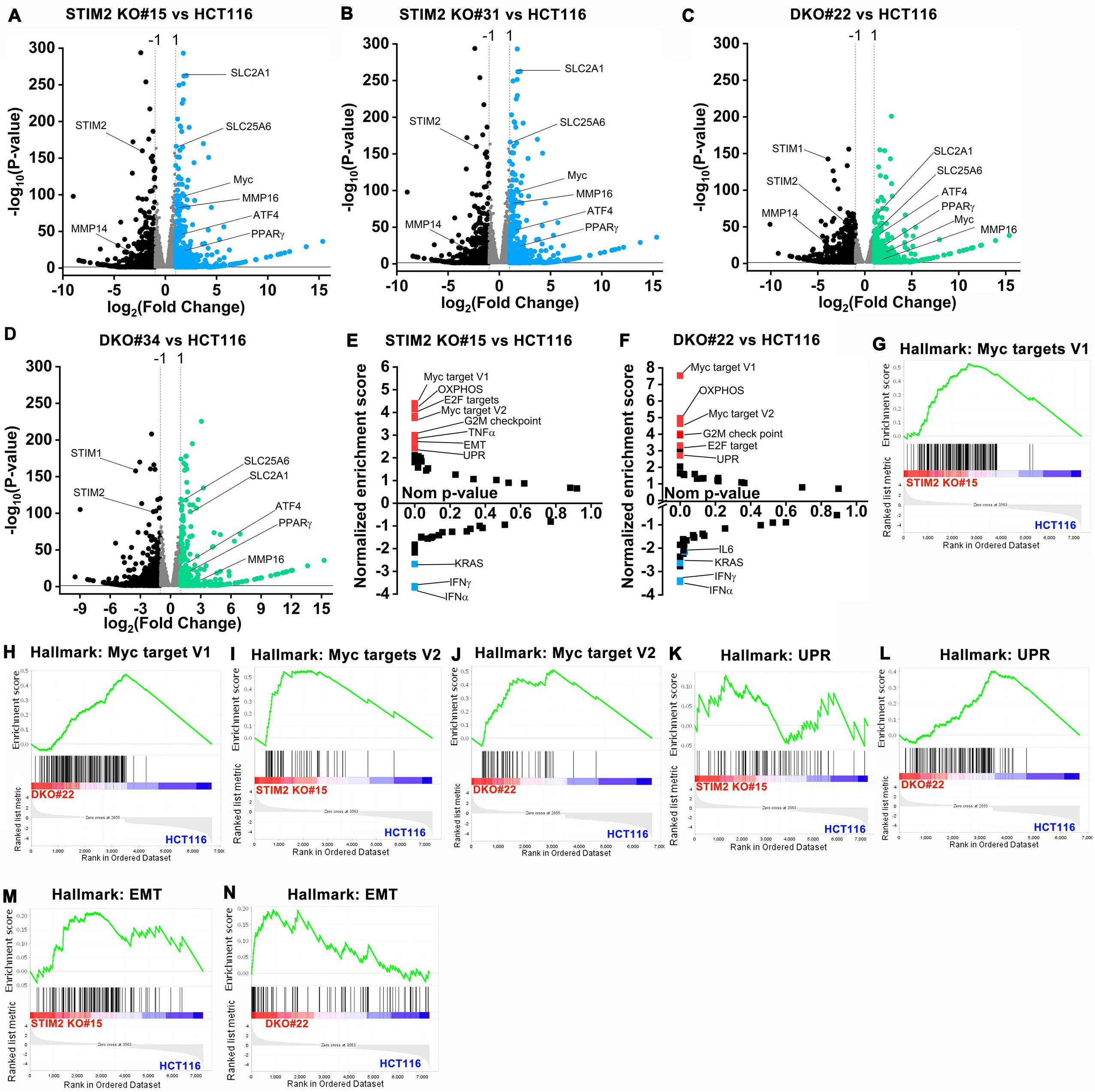
Loss of STIM2 leads to transcriptional reprogramming of CRC cells. (A-D) Volcano plot showing a comparison of differentially expressed genes between (A) STIM2 KO #15 and HCT116, (B) STIM2 KO #31 and HCT116, (C) DKO #22 and HCT116, and (D) DKO #34 and HCT116. The X-axis represents log_2_ fold change and Y axis -log_10_ (P-value). The threshold in the plot corresponds to P-value <0.05 and log 2-fold change <-1.0 or > 1.0. The significantly downregulated genes are represented in black and upregulated in blue or green. (E, F) The pathway analysis showing plot between normalized enrichment score and nominal p-value. The positive enrichment score represents upregulated pathway, and negative enrichment shows downregulated pathways in (E) STIM2KO #15 vs. HCT116 and (F) DKO #22 vs. HCT116. (G, H) GSEA analysis between (G) HCT116 and STIM2 KO #15 and (H) HCT116 and DKO #22 show a positive correlation in the enrichment of hallmark of Myc target version 1 genes. (I, J) GSEA analysis between (I) HCT116 and STIM2 KO #15 and (J) HCT116 and DKO #22 show a positive correlation in the enrichment of hallmark of Myc target version 2 genes. (K, L) GSEA analysis between (K) HCT116 and STIM2 KO #15 and (L) HCT116 and DKO #22 show a positive correlation in the enrichment of hallmark of unfolded protein response (UPR) genes. (M, N) GSEA analysis between (M) HCT116 and STIM2 KO #15 and (N) HCT116 and DKO #22 shows a positive correlation in the enrichment of hallmark of epithelial to mesenchymal transition (EMT) genes.

### Loss of STIM2 leads to increased SERCA2 and ER Ca^2+^ levels

It is well established that ER Ca^2+^ homeostasis is critical for ER function and protein synthesis and that altered ER Ca^2+^ levels are directly linked to ER stress and initiation of the unfolded protein response (UPR) (Corazzari *et al*., 2017; Mekahli *et al*, 2011; Preissler *et al*., 2020). Therefore, we performed direct ER Ca^2+^ measurement using the ER-targeted genetically encoded indicator, R- CEPIA1_ER_ (Suzuki *et al*, 2014). Cells were stimulated with 300 µM ATP in 0 mM Ca^2+^ in Hanks Balanced Salt Solution (HBSS) to cause maximal depletion of ER Ca^2+^, followed by ATP washout and addition of 2 mM Ca^2+^ to measure ER Ca^2+^ refilling. To standardize these experiments, we first transfected HCT116 and HEK293 cells with R-CEPIA1_ER_ and determined side by side ER Ca^2+^ release and refilling in these cells. Interestingly, the ER-Ca^2+^ depletion rate was significantly faster and refilling rate was slower in HCT116 cells compared the HEK293 cells **(Supplementary Figure 8 A-F)**. We performed similar experiments using STIM1 KO, STIM2 KO, and STIM DKO clones of HCT116 and DLD1 cells and we observed a significantly elevated basal ER Ca^2+^ in STIM2 KO and STIM DKO clones of HCT116 and DLD1 (**Figure 6 A-C and E-G**). The ER Ca^2+^ depletion rate was also increased in STIM2 KO and STIM DKO clones of HCT116 and DLD1 as compared to STIM1 KO clones and parental control cells (**Figure 6 D, H**), suggesting that overall ER Ca^2+^ levels were elevated in STIM2 KO and STIM DKO clones. We also measured ER Ca^2+^ levels in Orai1 KO and Orai TKO clones of HCT116. There was no marked difference observed in basal ER Ca^2+^ or ER Ca^2+^ depletion rate between Orai1 KO, Orai TKO cells compared to HCT116 control **(Supplementary Figure 8G-J)**. The Maximal ER Ca^2+^ refilling and ER Ca^2+^ refilling rate were significantly reduced in Orai1 KO and Orai TKO clones of HCT116 compared to control **(Supplementary Figure 8K, L)**. To determine the cause of increased ER Ca^2+^ in STIM2 KO and STIM DKO cells, we measured mRNA levels of SERCA. There was a significant increase in mRNA levels of SERCA2 in STIM2 KO and STIM DKO clones of HCT116 cells (**Figure 6I-K**). We also observed a marked increase in SERCA2 protein levels in STIM2 KO, and STIM DKO clones of HCT116 and DLD1 cells (**Figure 6L-Q**). SERCA2 protein levels were unaltered or slightly reduced in STIM1 KO clones of HCT116 and DLD1 cells (**Figure 6L-Q**). We measured the protein levels of the three IP_3_R subtypes and observed no change in protein levels between STIM1 KO, STIM2 KO, and STIM DKO clones of HCT116 and DLD1 cells **(Supplementary Figure 8M-S)**. Please note that the protein for the IP_3_R2 isoform was not detectable in HCT116 cells. Collectively, these data suggest that STIM2 regulates ER Ca^2+^ homeostasis, and loss of STIM2 increases basal ER Ca^2+^ by upregulating SERCA2 levels.

**Figure 6.**
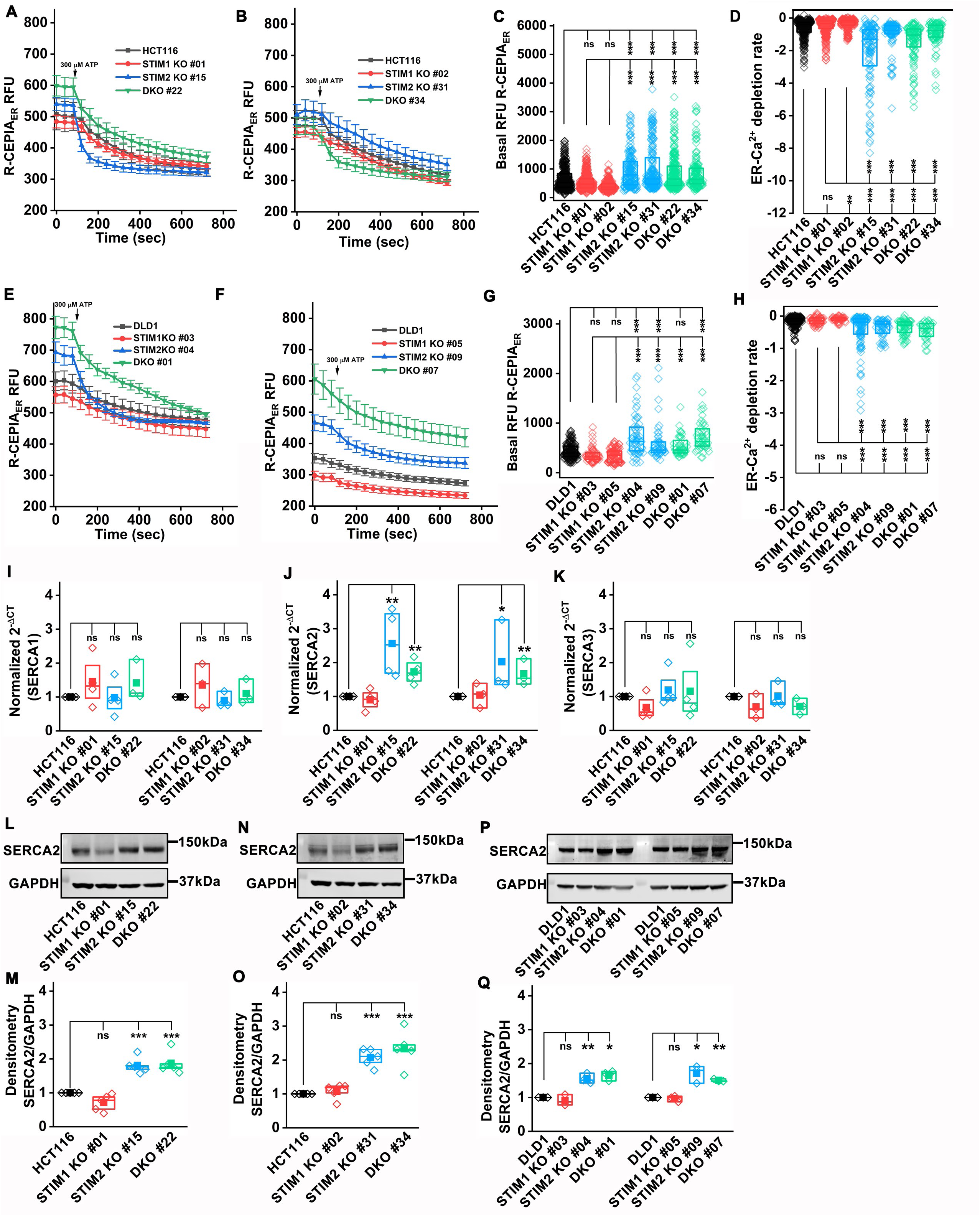
Loss of STIM2 leads to increased basal ER Ca^2+^. (A, B) ER Ca^2+^ measurement using genetically encoded R-CEPIA1_ER_. The ER Ca^2+^ depletion was stimulated with 300 µM ATP in 0 mM Ca^2+^ for 10 minutes. The graph represents the mean ± S.E.M. of (A) HCT116, STIM1 KO #01, STIM2 KO #15, and DKO #22 clone of HCT116, and (B) HCT116, STIM1 KO #02, STIM2 KO #31, DKO #34 clones of HCT116 cells. (C, D) Quantification of (C) Basal ER Ca^2+^, (D) ER Ca^2+^ depletion rate 3 min post stimulation in 0 Ca^2+^, HCT116 (n=350), STIM1 KO #01 (n=342), STIM1 KO#02 (n=210), STIM2 KO #15 (n=150), STIM2 KO #31 (n=170). DKO #22 (n=250), and DKO #34 (n=150) clones of HCT116. (E, F) ER Ca^2+^ measurement using genetically encoded R-CEPIA_ER_. The ER Ca^2+^ depletion was stimulated cells were stimulated with 300 µM ATP in 0 mM Ca^2+^ for 10 minutes. The graph represents the mean ± S.E.M. of (E) DLD1, STIM1 KO #03, STIM2 KO #04, and DKO #01 clone of DLD1, and (F) DLD1, STIM1 KO #05, STIM2 KO #09, DKO #07 clones of DLD1 cells. (G, H) Quantification of (G) Basal ER Ca^2+^, (H) ER Ca^2+^ depletion rate 3 min post stimulation in 0 Ca^2+^, in DLD1 (n=140), STIM1 KO #03 (n=54), STIM1 KO#05 (n=85), STIM2 KO #04 (n=50), STIM2 KO #09 (n=40). DKO #01 (n=35), and DKO #07 (n=40) clones of DLD1. (I-K) RT-qPCR analysis of mRNA levels of (I) SERCA1, (J) SERCA2, and (K) SERCA3 in HCT116, STIM1 KO, STIM2 KO, and DKO in HCT116 cells. (L-O) Measurement of SERCA2 protein level by probing the western blots with anti-SERCA2 and GAPDH antibody (L and N) representative blot probed and (M, and O) densitometric analysis of SERCA2 normalized to GAPDH in HCT116 and clones of STIM1 KO, STIM2 KO, and DKO in HCT116 cells. (P, Q) Measurement of SERCA2 protein level by probing the western blots with anti-SERCA2 and GAPDH antibody (P) representative blot, and (Q) densitometric analysis of SERCA2 normalized to GAPDH in DLD1 and clones of STIM1 KO, STIM2 KO, and DKO in DLD1 cells. All experiments were performed ≥three times with similar results. Statistical significance was calculated using one-way ANOVA followed by a post hoc Tukey test except for I-Q, where the paired t-test was used to compare between groups. *p<0.05, **p<0.01, ***p<0.001

### STIM2 deficiency promotes cMyc stabilization and enhances PPARɣ levels

The cMyc oncogene is a known target of chromosomal translocation and gene replication in cancer progression. cMyc is a transcription factor that induces proliferation by upregulating expression of genes that regulate the cell cycle, genes required for energy metabolism such as glycolytic genes, and ribosome biogenesis (Gao *et al*, 2009; Hart *et al*., 2012; Meyer & Penn, 2008). Ribosome biogenesis has been linked to increased protein synthesis regulated by cMyc activity and is required for survival of cancer cells (Arabi *et al*, 2005; Barna *et al*, 2008; Chan *et al*, 2011; Destefanis *et al*., 2020; Grandori *et al*, 2005; Iritani & Eisenman, 1999; Popay *et al*, 2021). RNA sequencing analysis of STIM2 KO and STIM DKO clones of HCT116, showed that cMyc targets and ribosome biogenesis genes pathways were enriched (**Figure 5, Supplementary Figure 6**), suggesting that cMyc is active in STIM2 KO and STIM DKO clones. Therefore, we measured cMyc levels in STIM1 KO, STIM2 KO, and STIM DKO clones of HCT116 and DLD1 cells using western blotting. Indeed, cMyc phosphorylation (Ser62) was markedly increased in both STIM2 KO and STIM DKO clones of HCT116 and DLD1 cells (**Figure 7 A, B**). There was no change in cMyc phosphorylation in STIM1 KO clones of HCT116 and DLD1, Orai1 KO, and Orai TKO clones of HCT116 cells as compared to their respective clones (**Figure 7A, B and Supplementary Figure 9A, B**). ERK is known to stabilize cMyc to regulate cancer cell proliferation and growth by regulating cell cycle checkpoint genes (Marampon *et al*, 2006; Sears *et al*, 2000; Tsai *et al*, 2012). STIM2 KO and STIM DKO clones of HCT116 and DLD1 had significantly increased ERK phosphorylation (Thr202/Tyr204) compared to their respective controls (**Figure 7A, C)**. ERK phosphorylation was unaltered in STIM1 KO, Orai1 KO, and Orai TKO clones as compared to control (**Figure 7A, C and Supplementary Figure 9C, D**). Although we did not see any significant changes in cMyc protein levels between wildtype and different STIM knockout clones of HCT116 and DLD1 cells, TCGA analysis using UALCAN showed that cMyc mRNA levels were significantly upregulated in primary tumors of COAD patients (**Supplementary Figure 9E**). The mRNA levels of cMyc were also upregulated in all cancer stages of COAD patients (**Supplementary Figure 9F**). Using cBioPortal, we found that cMyc expression has a stronger negative correlation with the expression of STIM2 than with that of STIM1 in CRC (**Supplementary Figure 9G, H**).

**Figure 7.**
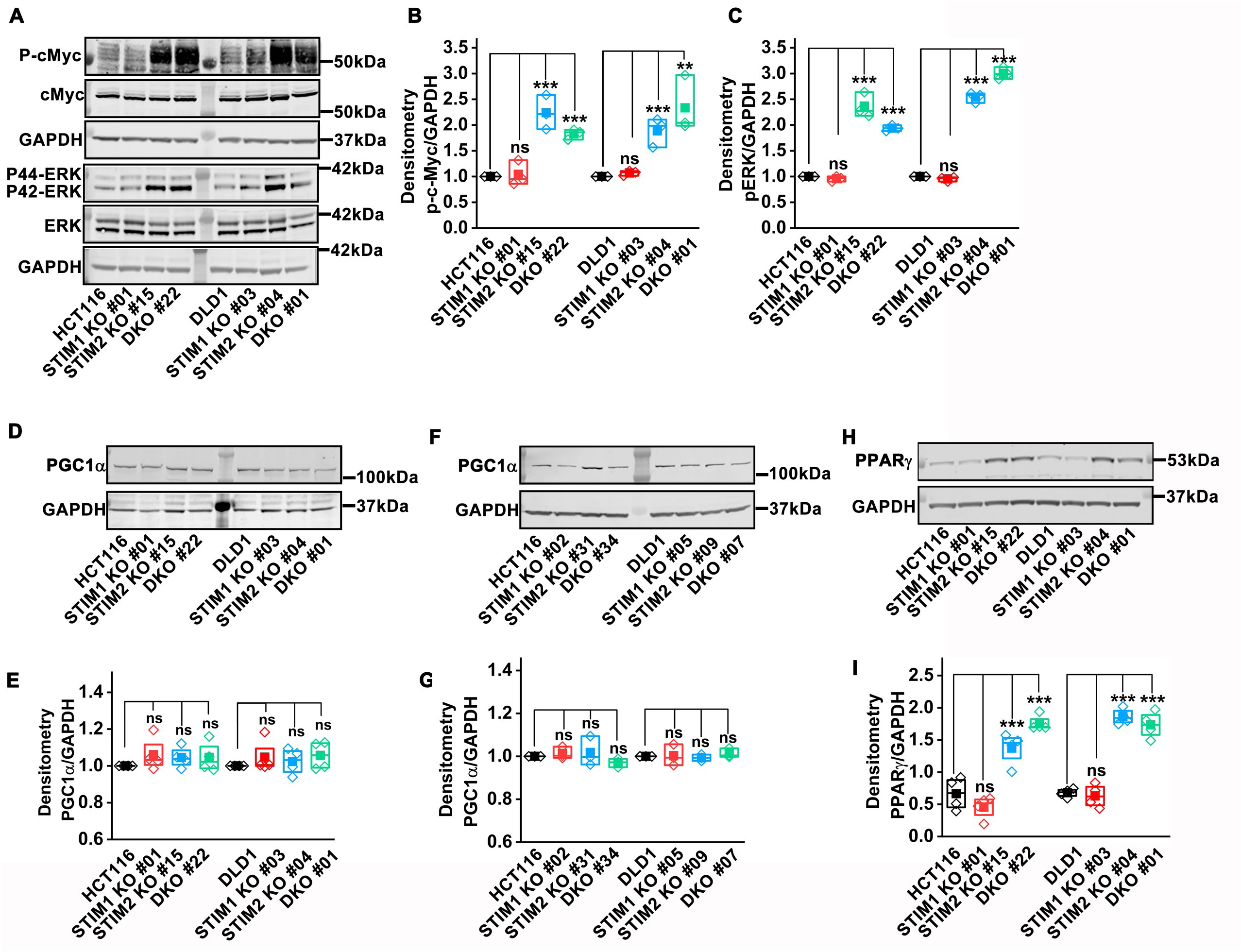
STIM2 deletion promotes cMyc stabilization and increases PPARɣ levels. (A) Representative western blot probed with anti-phospho-cMyc, cMyc, phospho-ERK, ERK, and GAPDH. (B, C) Densitometric analysis of (B) phospho-cMyc and (C) phospho-ERK in HCT116, STIM1 KO #01, STIM2 KO #15, DKO #22 clone in HCT116 and DLD1, STIM1 KO #03, STIM2 KO #04, DKO #01 clones in DLD1 cells. (D-G) PGC1α levels determined by western blot (D, F) blot probed with anti-PGC1α and GAPDH antibody and (E, G) densitometric analysis of PGC1α levels in HCT116, DLD1, STIM1 KO, STIM2 KO, and DKO clones of HCT116 and DLD1 cells. (H, I) PPARɣ levels quantified using (H) western blot probed with anti-PPARɣ and GAPDH antibody, and (I) densitometric analysis of PPARɣ levels in HCT116, DLD1, STIM1 KO, STIM2 KO, and DKO clones of HCT116 and DLD1 cells. All experiments were performed ≥three times with similar results. Statistical significance was calculated using paired t-test. *p<0.05, **p<0.01, ***p<0.001

We consistently observed that GLUT1 protein levels were significantly upregulated in STIM2 KO and STIM DKO clones of HCT116 and DLD1 cells (**Figure 4G, H, and Supplementary Figure 5C, D**). Since cMyc is a known molecular driver of glycolysis and tumor metabolic reprogramming (Vyas *et al*, 2016), we determined the role of STIM2 in regulating tumor metabolism through cMyc downstream targets. cMyc is known to regulate cellular energetics by affecting mitochondrial mass through the PGC1α/PPARɣ complex (Bost & Kaminski, 2019; Morrish & Hockenbery, 2014; Vyas *et al*., 2016). We observed a significant increase in mitochondrial biomass in both STIM2 KO and STIM DKO clones of HCT116 and DLD1 cells (**Figure 4I-Q**). We measured the protein levels of PGC1α and PPARɣ using western blotting. There was no significant change in PGC1α levels between STIM1 KO, STIM2 KO, STIM DKO, Orai1 KO and Orai TKO clones compared to control (**Figure 7D-G, and Supplementary Figure 9I, J**). However, PPARɣ levels were significantly upregulated in STIM2 KO and STIM DKO clones of HCT116 and DLD1 cells (**Figure 7H, I**). There was no apparent change in PPARɣ levels in STIM1 KO, Orai1 KO, and Orai TKO clones (**Figure 7H, I and Supplementary Figure 9K, L**), suggesting that loss of STIM2 leads to cMyc stabilization, increased PPARɣ and enhanced mitochondrial biogenesis to meet the energy demand of growing CRC cells.

### STIM2 deficiency promotes ATF4-dependent ER stress

The GSEA, KEGG, and DO analysis of the RNA sequencing data determined that cMyc targets, UPR, E2F targets, and ribosome biogenesis pathways were selectively enriched in STIM2 KO and STIM DKO clones of HCT116 (**Figure 5, Supplementary Figure 6**). Increased ribosome biogenesis is one of the indicators of increased protein synthesis, which is associated with UPR (Destefanis *et al*., 2020; Popay *et al*., 2021). We also observed a marked increase in SERCA2 protein levels and ER Ca^2+^ levels in STIM2 KO and STIM DKO cells (**Figure 6**). The disruption of ER Ca^2+^ homeostasis is known to induce ER stress (Corazzari *et al*., 2017; Deniaud *et al*, 2008; Mekahli *et al*., 2011). BiP is an ER chaperone, also known as GRP78/HSPA5, which regulates mammalian ER stress response. BiP levels were significantly and selectively upregulated in STIM2 KO and STIM DKO clones of HCT116 and DLD1 cells (**Figure 8A, B**). No apparent increase was observed in BiP levels in STIM1 KO clones of HCT116 and DLD1, and in Orai1 KO and Orai TKO clones of HCT116 as compared to their respective controls (**Figure 8A, B and Supplementary Figure 10G, H**). To determine which branch of ER stress pathway is triggered by the loss of STIM2, we measured ATF4, IRE1α and ATF6 levels and found that only ATF4 levels were significantly upregulated in STIM2 KO and STIM DKO clones of HCT116 and DLD1 (**Figure 8C, D and Supplementary Figure 10A, B**). There was no change in IRE1α levels in STIM1 KO, STIM2 KO, and STIM DKO clones of HCT116 and DLD1 cells (**Figure 8E, F and Supplementary Figure 10C, D**). Except for DLD1 STIM2 KO #04 cells, we did not observe a change in ATF6 levels in STIM1 KO, STIM2 KO, and STIM DKO clones of HCT116 and DLD1 cells **(Figure 8G, H and Supplementary Figure 10E, F)**. Furthermore, Orai1 KO and Orai TKO clones of HCT116 did not show any changes in ATF4 and IRE1α levels (**Supplementary Figure 10I-L**). Surprisingly, ATF6 levels were significantly upregulated in Orai1 KO clones but not in Orai TKO clones of HCT116 (**Supplementary Figure 10M, N**). We also determined PERK and eIF2α levels and found that the phosphorylation of eIF2α was significantly upregulated in STIM2 KO and STIM DKO clones of HCT116 cells (**Figure 8I, J**). PERK phosphorylation showed strong variability and while PERK phosphorylation trended towards an increase in STIM2 KO and STIM DKO cells, this increase did not reach statistical significance (**Figure 8K, L**). These data show that the BiP/eIF2α/ATF4 pathway is activated in response to ER stress in STIM2 KO and STIM DKO clones of HCT116 and DLD1 cells. This ER stress pathway is not active in STIM1 KO, Orai1 KO, and Orai TKO cells, further supporting that STIM2 regulates ER stress in a SOCE-independent manner.

**Figure 8.**
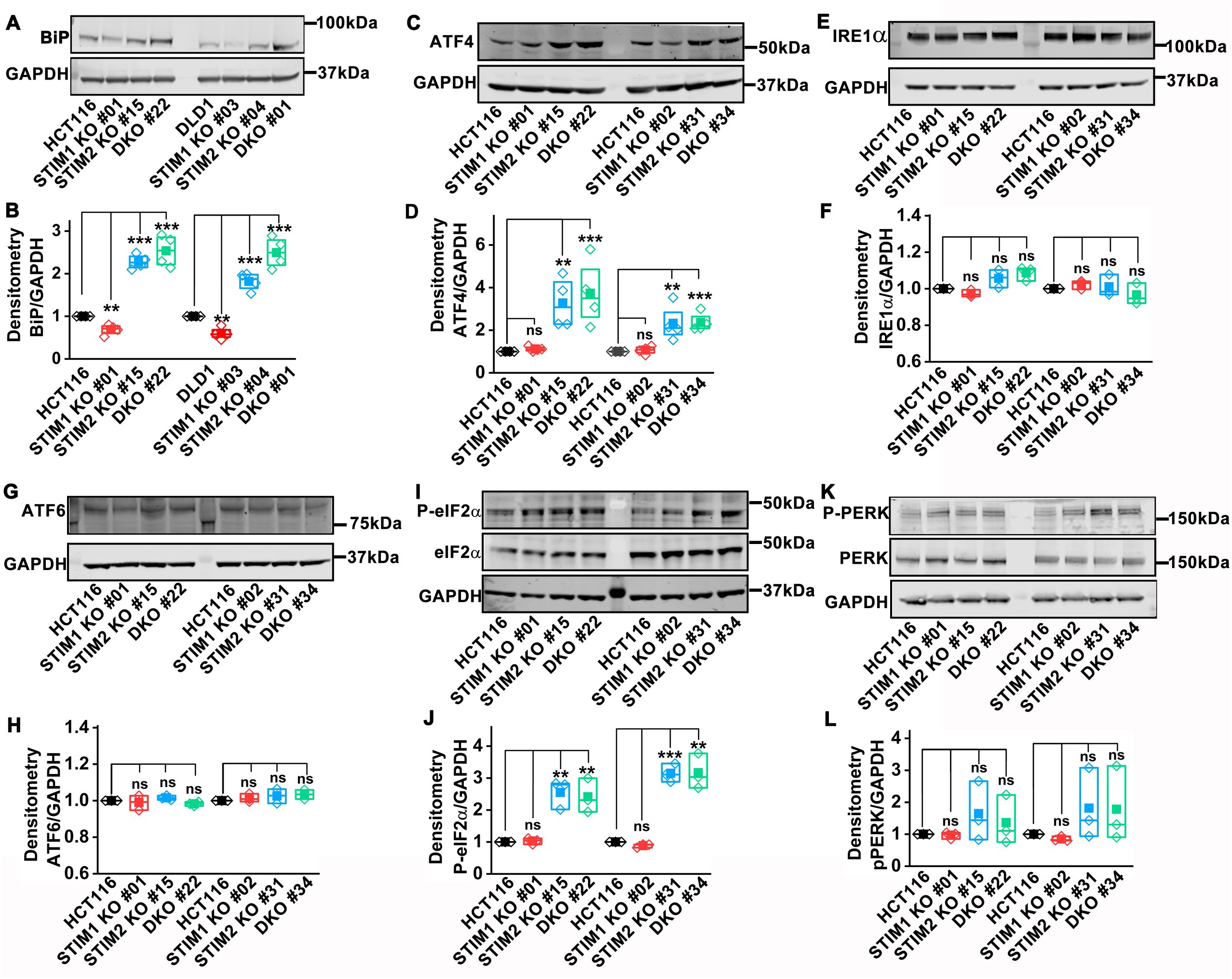
Loss of STIM2 activates the ATF4-dependent ER stress pathway. (A, B) Quantification of BiP protein levels using (A) western blot probed with anti-Bip and GAPDH antibody, and (B) densitometric analysis in HCT116, DLD1, STIM1 KO, STIM2 KO, and DKO clones of HCT116 and DLD1cells. (C, D) Quantification of ATF4 protein levels using (C) western blot probed with anti-ATF4 and GAPDH antibody, and (D) densitometric analysis in HCT116, STIM1 KO, STIM2 KO, and DKO clones of HCT116. (E, F) Quantification of IRE1α protein levels using (E) western blot probed with anti-IRE1α and GAPDH antibody, and (F) densitometric analysis in HCT116, STIM1 KO, STIM2 KO, and DKO clones of HCT116. (G, H) Quantification of ATF6 protein levels using (G) western blot probed with anti-ATF6 and GAPDH antibody, and (H) densitometric analysis in HCT116, STIM1 KO, STIM2 KO, and DKO clones of HCT116. (I, J) Quantification of phospho-eIF2α protein levels using (I) western blot probed with anti-phospho-eIF2α, eIF2α, and GAPDH antibody, and (J) densitometric analysis in HCT116, STIM1 KO, STIM2 KO, and DKO clones of HCT116. (K, L) Quantification of phospho-PERK protein levels using (K) western blot probed with anti-phospho-PERK, PERK, and GAPDH antibody, and (L) densitometric analysis in HCT116, STIM1 KO, STIM2 KO, and DKO clones of HCT116. All experiments were performed ≥three times with similar results. Statistical significance was calculated using paired t-test. *p<0.05, **p<0.01, ***p<0.001

## Discussion

Ca^2+^ ions play a critical role in a myriad of biological functions, including cellular metabolism, bioenergetics, transcription, cellular proliferation, cell death, and cellular stress responses. Recent advances have shown that compartmentalized Ca^2+^ within the ER, mitochondria, endolysosomes and Golgi are interconnectedly and tightly regulated and this regulation is essential for normal cellular function (Stewart *et al*, 2015). Perturbed Ca^2+^ homeostasis within cancer cells alters their transcriptional profile to support requirements for growth and metastasis (Jones & Hazlehurst, 2021; Marchi & Pinton, 2016; Pathak & Trebak, 2018; Zheng *et al*, 2022). Mitochondria and ER are tightly interlinked, and perturbation of Ca^2+^ homeostasis of any of these organelles strongly affects their function (Bustos *et al*, 2017). Changes in mitochondrial Ca^2+^ homeostasis have been implicated in multiple disease conditions, including cancer (Pathak & Trebak, 2018; Vasan *et al*, 2020). ER Ca^2+^ regulates protein synthesis, ER structure, function, and ER stress responses (Moy *et al*, 2022; Papp *et al*, 2012). Multiple cancer types have been linked to ER stress, which regulates cancer progression through transcriptional reprograming and metabolic transformation to meet the needs for cancer growth and dissemination (Chen & Cubillos-Ruiz, 2021; Corazzari *et al*., 2017).

Even though STIM2 is one of the major components of SOCE, its role in cancer progression has remained obscure. Our TCGA data analysis of colorectal cancer patients showed that STIM2 mRNA levels are unaltered while STIM1 mRNA levels were significantly reduced in all stages of cancer progression. However, survival analysis of these patients showed a strong correlation between low STIM2 mRNA levels and poor prognosis in both COAD and READ patients. Furthermore, we found no correlation between STIM1, Orai1 or Orai2 expression and survival of COAD and READ patients. Interestingly, Orai3 levels were significantly upregulated in primary tumors, and higher Orai3 mRNA level was strongly correlated with poor prognosis of CRC patients. As such, future studies are required to determine the function of Orai3 in CRC progression.

In the past few years, many studies have shown a pro-growth function of STIM1 and Orai1 in various cancer types, including breast cancer, colorectal cancer, glioblastoma, and melanoma (Khan *et al*, 2020; Mo & Yang, 2018; Motiani *et al*, 2010; Motiani *et al*, 2013a; Motiani *et al*, 2013b; Shapovalov *et al*, 2021; Xie *et al*, 2016). In most cancer cell types, blocking or reducing STIM1/Orai1 function led to reduced SOCE and subsequent reduction in proliferation and migration (Faris *et al*, 2022; Motiani *et al*., 2013a; Moy *et al*., 2022; Wang *et al*, 2015; Yang *et al*, 2013; Yang *et al*, 2009; Zhang *et al*, 2015). Consistently, we show here that loss of STIM, or Orai proteins leads to reduced proliferation. When we measured colorectal cancer cell migration using the wound healing assay, we saw significantly increased migration of cells lacking STIM or Orai proteins. However, serum-induced migration and invasion in Boyden chamber assays showed reduced migration and invasion of STIM1 KO cells while migration and invasion of STIM2 KO and STIM DKO cells were greatly increased. Cell migration in wound healing assays is mostly driven by cell-cell interactions while migration in Boyden chamber assays is largely contributed by chemotaxis. Consequently, we used spheroid to assess proliferation, migration, and invasion and found that the spheroid size was significantly larger in STIM2 KO and STIM DKO cells. The spheroids of STIM2 KO and STIM DKO cells also had a significant amount of loosely attached cells, indicating a pro-migratory and pro-metastatic phenotype. Surprisingly, STIM DKO cells showed a phenotype similar to that of STIM2 KO cells rather than STIM1 KO, strongly arguing that the phenotype of STIM2 KO cells is Orai- and SOCE-independent. Our *in vivo* xenograft studies showed enhanced tumor growth and metastasis specifically in STIM2 KO and STIM DKO cells. Interestingly, Aytes et al. reported that STIM2 overexpression in colorectal cancer cells confers a tumor suppressor phenotype (Khan *et al*., 2020; Wang *et al*., 2015; Yang *et al*., 2009). Although these studies did not provide a mechanism by which STIM2 suppresses tumor growth, they agree with our findings.

While reduced SOCE by loss of STIM1 or Orai1 in normal cells reduces mitochondrial metabolism and glycolysis (Emrich *et al*., 2023; Johnson *et al*., 2022; Maus *et al*, 2017; Novakovic *et al*., 2023; Vaeth *et al*., 2017; Yoast *et al*, 2021), the function of STIM2 in cellular metabolism of either normal or cancerous cells has remained unclear. Here we show that loss of STIM2 results in increased glycolysis, mitochondrial respiration, and mitochondrial biogenesis. Similarly, STIM1/2 double knockout cells showed a similar phenotype. However, Orai1 KO and Orai1/2/3 triple KO cells behaved in a similar fashion to STIM1 KO cells, where glycolysis and mitochondrial respiration were reduced. These findings underscore that STIM2 drives the metabolic transformation of STIM2 KO and STIM DKO cells in a SOCE-independent manner. Further, the activity of cMyc, a known regulator of transcription of genes required for metabolic transformation, proliferation, and metastasis (Li *et al*., 2005; Morrish & Hockenbery, 2014; Satoh *et al*., 2017), was significantly enhanced in both STIM2 KO and STIM DKO cells and cMyc expression negatively correlates with STIM2 expression in CRC patients, suggesting that CRC tumors lose STIM2 to support an optimized ER stress response and cMyc-dependent metabolic reprogramming. Indeed, Tameire et al showed that cMyc activity upregulates ATF4 to relieve cMyc-dependent proteotoxic stress and balance between enhanced protein translation and cancer cell survival/tumor progression. (Tameire *et al*., 2019). The PPARɣ, a nuclear receptor and known regulator of mitochondrial metabolism genes, was also upregulated in STIM2 KO and STIM DKO cells, suggesting that cMyc might be increasing PPARɣ levels to enhance mitochondrial biogenesis. However, further studies are required to understand how STIM2 loss regulates cMyc/ PPARɣ levels in CRC cells.

The canonical CRC subtype called consensus molecular subtype 2 (CMS2) is characterized by the strong upregulation of cMyc downstream signaling while the CMS3 subtype is known for the enrichment of multiple metabolism signatures (Guinney *et al*., 2015). The loss of STIM2 caused marked changes in the transcriptional profile of CRC cells, and this change in transcriptional profile matches both canonical CRC subtypes, CMS2 and CMS3. This suggests that loss of STIM2 is likely correlated with poor prognosis in both CMS2 and CMS3 CRC subtypes. Subtype probability scoring determined that CMS2 and CMS3 are closely related and hard to disambiguate (Guinney *et al*., 2015), in further support of the idea that STIM2 regulates both CMS2 and CMS3 CRC subtypes.

STIM1 and Orai1 are the major regulators of SOCE, and their activation has been studied extensively (Lewis, 2020; Yeung *et al*, 2020). Multiple studies show that under basal conditions and upon moderate ER Ca^2+^ depletion, STIM2 activates SOCE (Brandman *et al*, 2007a; Emrich *et al*., 2021). In HCT116 and DLD1 CRC cells, maximal activation with 300 µM ATP revealed a complete loss of SOCE in STIM1 KO and STIM DKO conditions with ∼ 50% loss of SOCE in STIM2 KO cells. However, both *in vivo* and *in vitro* CRC progression phenotypes were the same between STIM2 KO and STIM DKO cells, suggesting that STIM2 functions in a SOCE-independent mechanism to regulate CRC progression. Our direct measurements of ER Ca^2+^ levels using R-CEPIA_ER_ showed that basal ER Ca^2+^ and SERCA2 protein levels were significantly enhanced in STIM2 KO and STIM DKO cells. Perturbation of ER Ca^2+^ is known to alter mitochondrial Ca^2+^ homeostasis and subsequently cause mitochondrial stress and alter metabolism (Bustos *et al*., 2017; Marchi & Pinton, 2016). Although, it is well established that ER Ca^2+^ depletion activates ER stress responses (Corazzari *et al*., 2017; Krebs *et al*, 2015; Zheng *et al*., 2022), our data indicate that increased ER Ca^2+^ is also associated with selective activation of the ATF4 branch of ER stress. Our RNA sequencing analysis showed that ribosome biogenesis is selectively increased in STIM2 KO and STIM DKO CRC cells, which is a hallmark of increased protein synthesis. We showed that the eIF2α and ATF4 levels are upregulated in STIM2 KO and STIM DKO CRC cells, Therefore, our data suggest that loss of STIM2 initiates the activation of the ATF4 branch of ER stress, likely in response to enhanced ER Ca^2+^. However, how enhanced ER Ca^2+^ contributes to ER stress and the mechanisms involved require further investigations.

In summary, we have identified a SOCE-independent role of STIM2 in colorectal cancer progression, whereby the loss of STIM2 leads to enhanced tumor growth, metastasis, and poor prognosis. Mechanistically, loss of STIM2 causes increased ER Ca^2+^ levels, protein synthesis, and cMyc stabilization, leading to activation of ER stress response in ATF4 dependent manner. The activation of the ATF4 ER stress pathway causes transcriptional reprogramming and metabolic transformation to support increased metastasis of CRC cells. These changes are the markers of CMS2, and CMS3 subtypes of CRC. Therefore, STIM2 could serve in CRC prognosis or in targeted therapies to inhibit CRC tumor growth and metastasis.

## Methods

Key resource table:

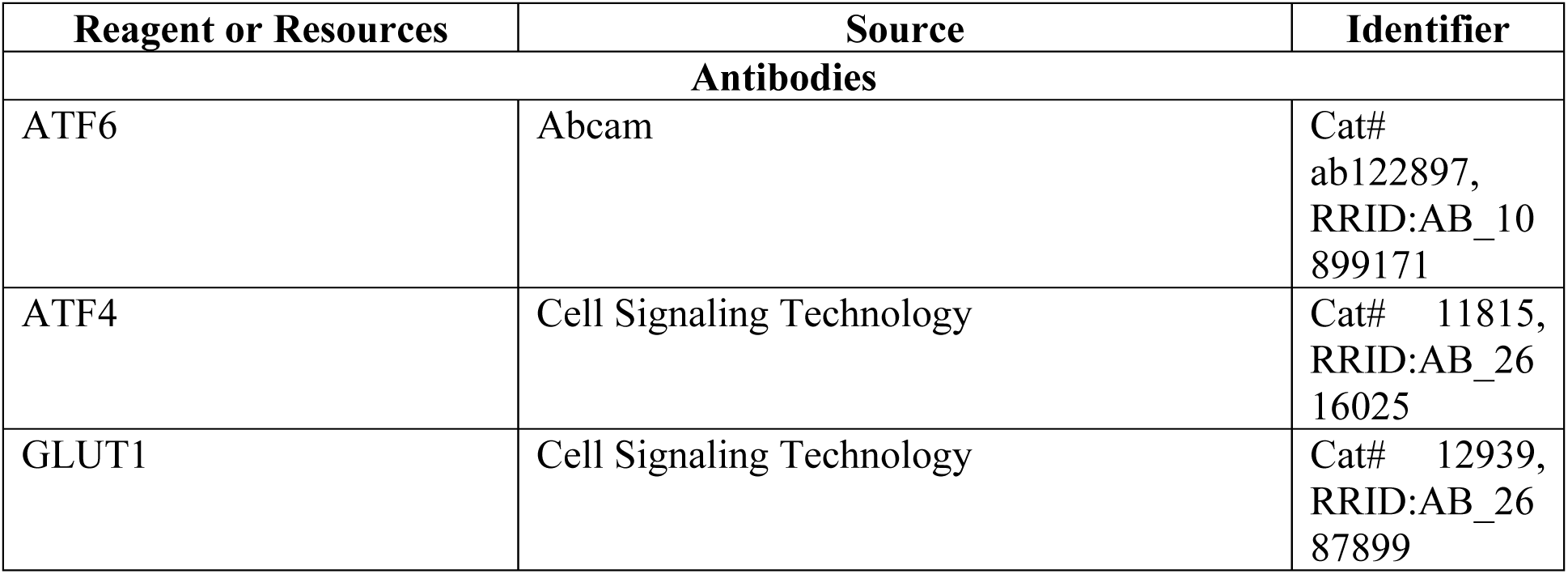
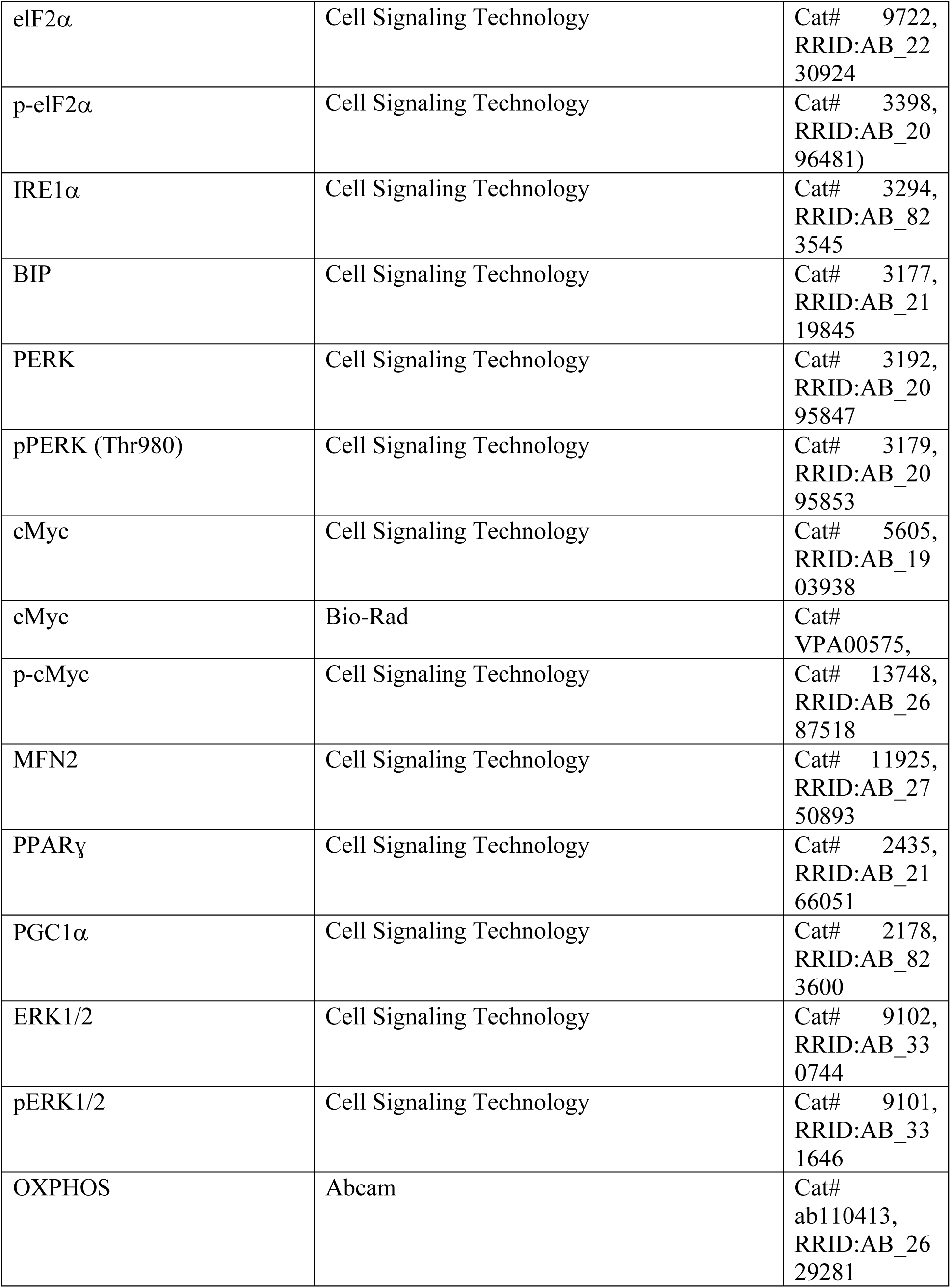
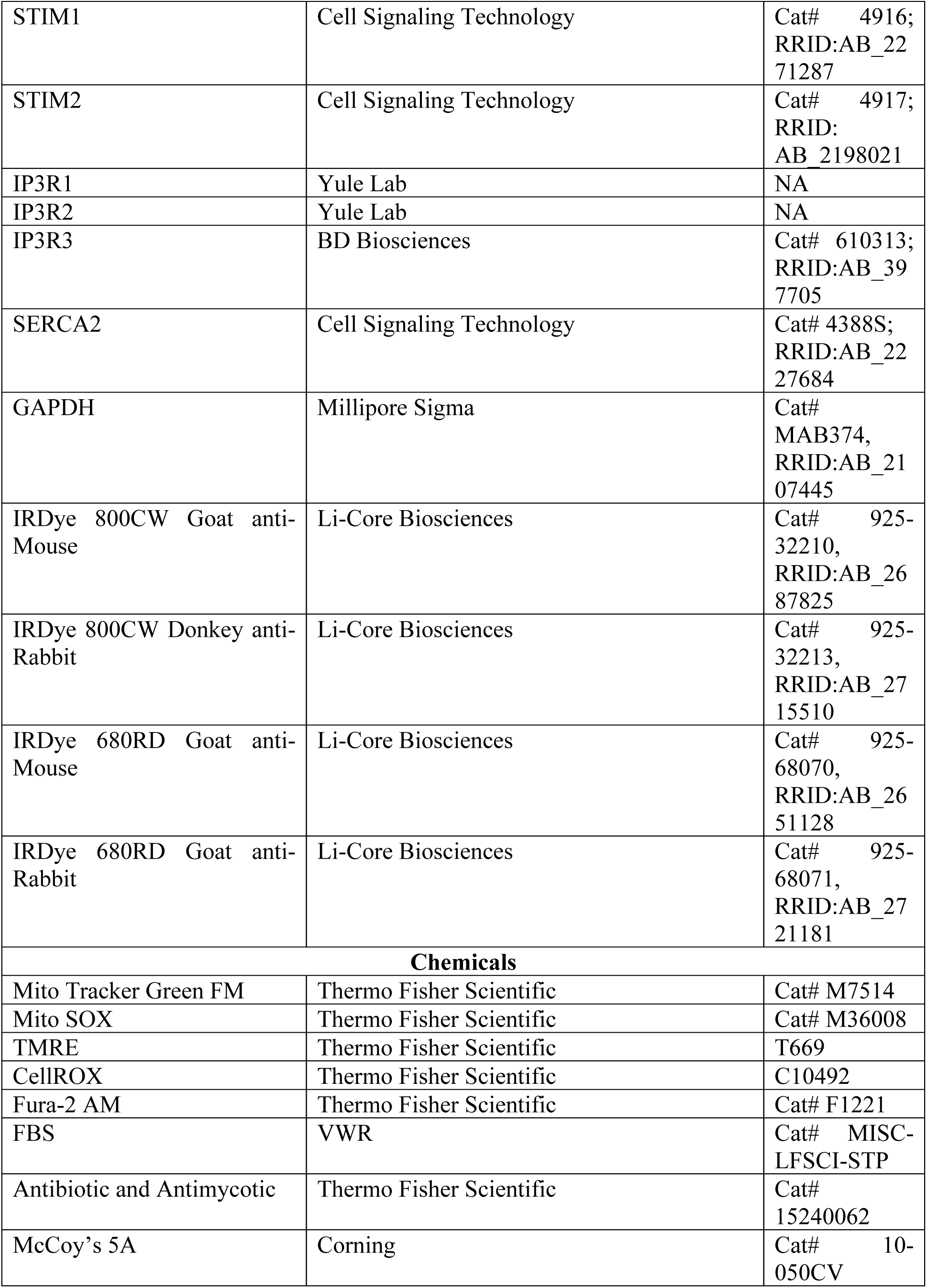
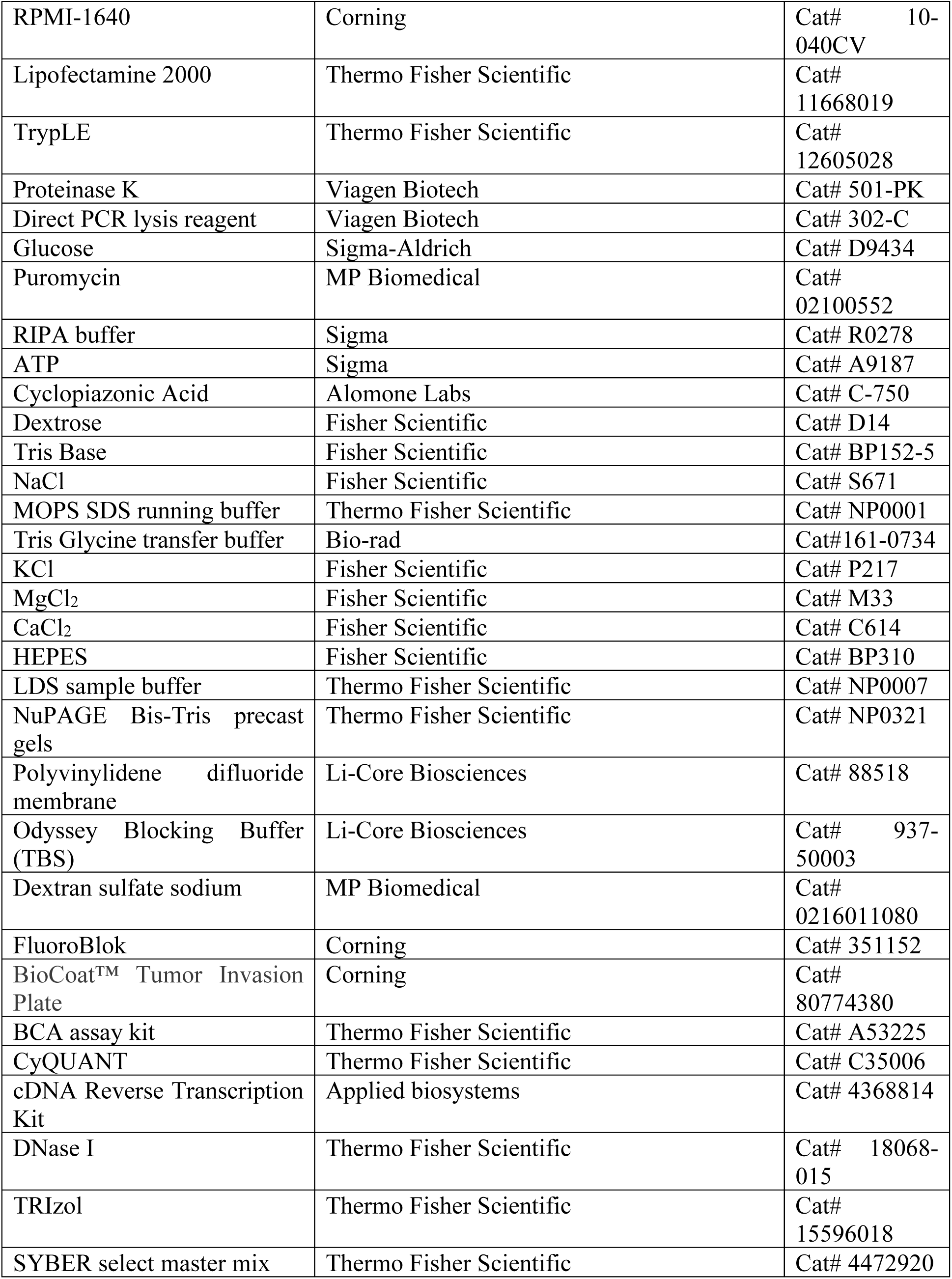
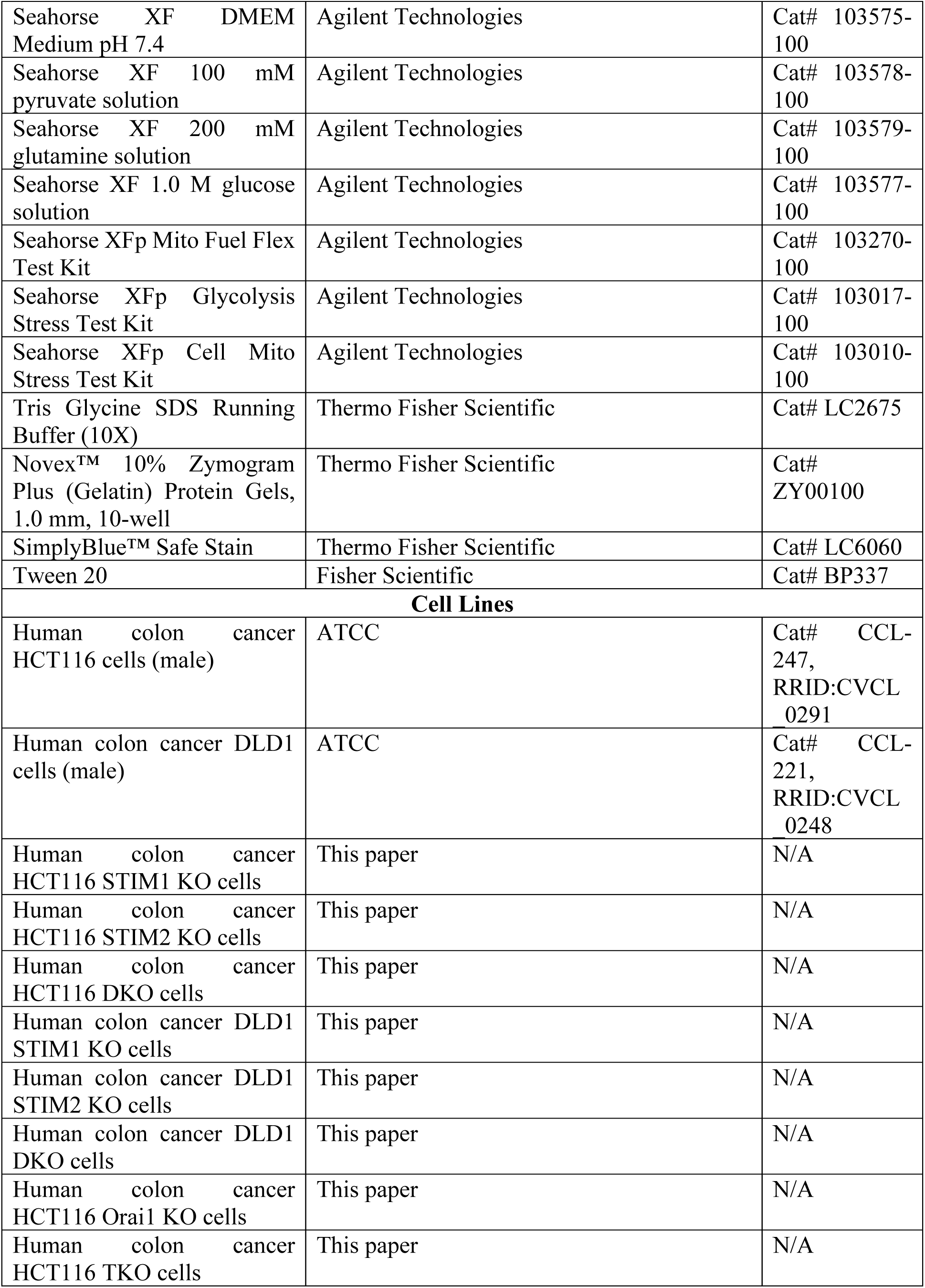
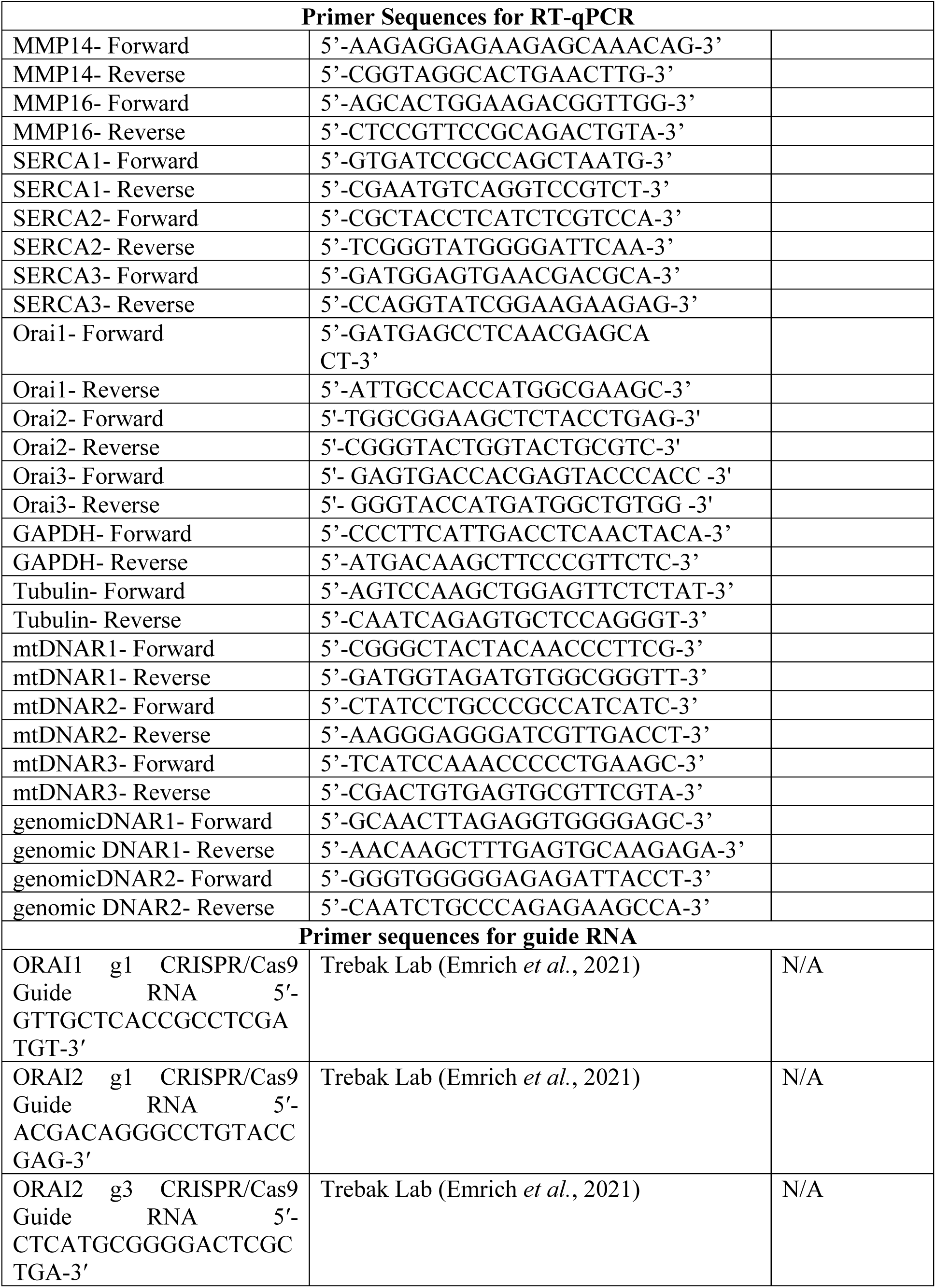
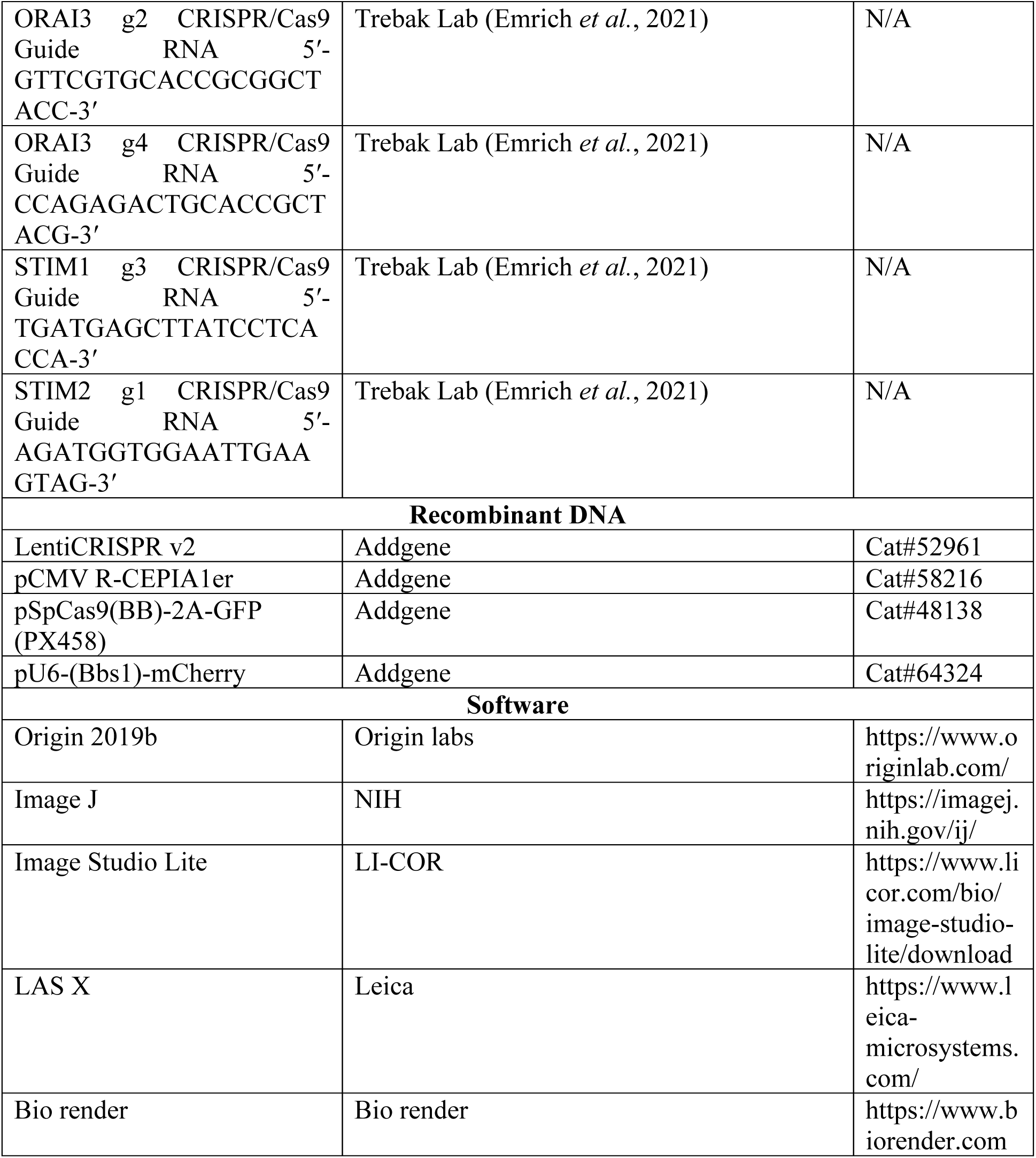

### Mice

NOD SCID (NOD.CB17-*Prkdc^scid^*/J) were also obtained from the Jackson laboratories. The mice were housed in 12:12 hr light dark cycle (22°C to 25°C) in an aseptic environment at The Pennsylvania State University animal facility. The mice were euthanized at the end of the experiment. The tumors and organs were collected for histopathology, protein, and mRNA analysis. All the mouse experiments were conducted in accordance with The Pennsylvania State University Institutional Animal Care and Research Advisory Committee.

#### Cell culture and plasmid transfection

HCT116 and DLD1 cells were cultured in McCoy’s 5A and RPMI-1640 media respectively and supplemented with 10% FBS and 1X antibiotic-antimycotic (AA) agent. All the cells were kept at 37°C in 5% CO_2_ incubator. For the transfection, the cells were cultured at 60-70% confluency. Next day, the cells were transfected with plasmid using Lipofectamine 2000 reagent, according to manufacturer protocol (Invitrogen).

### Plasmids, CRISPR/Cas9, and Lentiviral infection

Briefly, at the time of transfection the cells were seeded at 60-70% confluency. The plasmid, and Lipofectamine 2000 were diluted in Opti-MEM and mixed. The mixture was incubated for 5-10 min at room temperature, and then added to the cells cultured in media without AA. The medium of transfected cells was changed to normal medium media with 10% FBS and 1X AA after 4-6 hr. The expression of plasmid was confirmed 24 to 72 hr after transfection.

We generated several clones of STIM1, STIM2 and STIM1/2 DKO knockout in both HCT116 and DLD1 cells using the CRISPR/Cas9 system as described previously (Emrich *et al*., 2021; Pathak *et al*., 2020; Yoast *et al*, 2020b).Briefly, highly sequence specific guide RNAs (gRNA) targeting STIM1 and STIM2 were cloned in either pSpCas9(BB)-2A-GFP or pU6-(BbsI)_CBh-Cas9-T2A-mCherry vector. Single gRNA was used to generate STIM1 or STIM2 KO cells. STIM2 KO cells were transfected with STIM1 gRNA to generate STIM1/2 double knockout cells. The cells transfected with gRNA were individually selected for the mChery/GFP markers by FACS (BD FACS Aria SORP) in 96 well plate. To reduce potential off-target effects of the CRISPR/Cas9 system, we generated several independent clones of each knock out cell line.

### Transcriptome sequencing (RNA-seq) and analysis

To perform RNA-seq mRNA was isolated from total RNA (minimum 3 µg) from HCT116, STIM1 KO, STIM2 KO, and DKO HCT116 cells using the RNeasy Mini Kit. Short fragment off mRNA was generated using fragmentation buffer (Ambion). The double-stranded cDNA was generated, end-repaired, and ligated to Illumina adapters. These cDNAs were size selected (∼250 bp) and PCR amplified. The cDNA library was fed to the Illumina HiSeq 2500 sequencing platform (Berry Genomics) and the expression level of each transcript of a gene was quantified using read counts with HTseq. The genes with identifiers that has zero read counts were removed from each sample, and the biomaRt R package (Durinck *et al*, 2009) was used to convert Ensemble gene identifiers to gene symbols using the January, 2019 archive (http://jan2019.archive.ensembl.org/). Identifiers corresponding to multiple gene symbols were removed, as they were lowly expressed genes.

Exploratory analyses were performed after utilizing the VST (variance stabilizing transformation) function in the DESeq2 R package (Love *et al*, 2014) to apply a variance stabilizing transformation to the read count data. The results strongly suggested the presence of a batch effect. Therefore, the design matrix used in the differential expression analysis included batch as a covariate. After applying the voom transformation (Law *et al*, 2014) to the read counts, the limma R package (Ritchie *et al*, 2015) was used to identify differentially expressed genes based on a q-value threshold of 0.05. The RNA seq data are publicly available in NCBI with BioProject number PRJNA984240.

### Gene set enrichment analysis (GSEA)

The gene set enrichment analysis (GSEA) software (Subramanian *et al*, 2005) was used to perform pathway analyses using the limma output. Briefly, genes in the limma output were ordered according to the *t* test statistic. Then a GSEA pre-ranked analysis was performed using the Hallmark Gene Sets in the Molecular Signatures Database (http://software.broadinstitute.org/gsea/msigdb/index.jsp).

### TCGA analysis

RNA-sequencing (RNA-seq) based gene expression data for both tumor and normal samples and clinical data for the combined TCGA colon adenocarcinoma and rectal adenocarcinoma (COADREAD) cohort was downloaded from the Broad Institute’s Firehose GDAC (https://gdac.broadinstitute.org/). Gene expression was quantified as log2 (normalized RSEM+1). Somatic gene mutation data was obtained from (Ellrott *et al*, 2018) after restricting to samples in the RNA-seq cohort. Kruskal-Wallis and Wilcoxon rank sum tests and tests were used to compare expression values across groups defined by the clinical and mutation data.

### Quantitative Reverse Transcription-PCR (qPCR)

Briefly the cells were washed with cold PBS and RNA was isolated using RNeasy Mini Kit. High-Capacity cDNA Reverse Transcription Kit (Applied Biosystems, Foster City, CA) was used to make cDNA and the cDNA was further used for quantitative real-time PCR using SYBR select master mix (Thermo Scientific, Wilmington, DE, USA) according to the manufacturer protocol. The reactions were run in technical duplicates using Quant Studio 3 system (Applied Biosystems, Foster City, CA). The expression levels of genes were normalized to at least two housekeeping gens, GAPDH or Tubulin (mentioned in the figure legends) using 2^-ΔCt^ method.

### Western blot

As described previously (Pathak *et al*., 2020), briefly the cells were cultured and after attaining 80-90% confluency the cells were lysed in 100 µl RIPA buffer (Sigma) containing protease and phosphatase Inhibitor. 30-50 µg of the protein was loaded on an 4-12% gel (NuPAGE Bis-Tris precast gels, Life Technologies) and transferred to an activated polyvinylidene difluoride membrane. The membrane was incubated in the Odyssey blocking buffer for 1 hr at room temperature and then incubated overnight at 4°C in primary antibody (1:1000) diluted in Odyssey blocking buffer. Next day, the membrane was washed with 0.1% TBST for 5 min (three times) at room temperature. The blot was incubation in IRDye for 1 hour at room temperature. The IRDyes used are mentioned in the key resource table and they were diluted in Odyssey blocking buffer (IRDye 680RD Goat anti-Mouse at 1:10,000 dilution and IRDye 800CW Donkey anti-Rabbit at 1:5000). The membrane was washed with 0.1% TBST for 5 minutes (three times) at room temperature and imaged in Odyssey CLx Imaging System (LI-COR, NE, USA). Densiometric analysis of the bands on membrane was performed using ImageJ.

### Metabolic profiling

The metabolic profiling of HCT116, STIM1 KO, STIM2 KO, and DKO HCT116 cells was performed using LC/MS. The LC/MS analyses were performed using Q Exactive Hybrid Quadrupole Orbitrap mass spectrometer. Which was equipped with an Ion Max source and a HESI II probe coupled to a Dionex UltiMate 3000 UPLC system (Thermo Fisher Scientific, San Jose, CA). The standard calibration mixture was used for external mass calibration and the calibration was performed every 7 days. The HCT116 and HCT116, STIM1 KO, STIM2 KO, and DKO HCT116 cells were collected by scrapping and cell pellets were washed with chilled PBS. Chilled LC/MS grade methanol (600 µl), water (300µl), and chloroform (400 µl) with 10 ng/ml valine-d8 as an internal standard was added to the cell pellet and vortexed for 10 min. The mixture was centrifuged for 10 min at 4°Cat max speed. Carefully, two layers were collected, upper layer for polar and bottom layer for non-polar metabolites. Both the samples were dried on table-top vacuum centrifuge and stored at −80°C. The dried samples were then resuspended in 100 mL water; 1 mL of each sample was injected onto a ZIC-pHILIC 2.1 3 150 mm (5 mm particle size) column (EMD Millipore). Buffer A was 20 mM ammonium carbonate, 0.1% ammonium hydroxide; buffer B was acetonitrile. The chromatographic gradient was run at a flow rate of 0.150 ml/min as follows: 0– 20 min.: linear gradient from 80% to 20% B; 20–20.5 min.: linear gradient from 20% to 80% B; 20.5–28 min.: hold at 80% B. The mass spectrometer was operated in full-scan, polarity switching mode with the spray voltage set to 3.0 kV, the heated capillary held at 275°C, and the HESI probe held at 350°C. The sheath gas flow was set to 40 units, the auxiliary gas flow was set to 15 units, and the sweep gas flow was set to 1 unit. The MS data acquisition was performed in a range of 70–1000 m/z, with the resolution set at 70,000, the AGC target at 106, and the maximum injection time at 80 msec. Relative quantitation of polar metabolites was performed with XCalibur QuanBrowser 2.2 (Thermo Fisher Scientific) using a 5 ppm mass tolerance and referencing an in-house library of chemical standards. Relative abundance of metabolites was calculated by normalizing profiling results to cell number.

### Mouse tumor xenograft experiments

Mice were anesthetized using isoflurane and after proper sterile preparation of the abdomen, a small (4-8 mm) incision was made by sharp sterile blade over the left upper quadrant of the abdomen. The peritoneal cavity was carefully exposed, and the spleen was located. Further, the spleen was carefully exteriorized on the sterile field around the incision area and 500,000 cells suspended in 50 ul McCoy’s (10 or 20% FBS) were then injected into the spleen via a 1 ml insulin syringe. A small bleb was observed while injecting the cells in the spleen confirming that the cells were successfully injected. The injection area was wiped with sterile cotton swab. The spleen was carefully placed back into the peritoneal cavity. Both peritoneal cavity and skin were closed with sterile absorbable suture.

### Intracellular Ca^2+^ measurements

Intracellular Ca^2+^ imaging was performed as described previously (Emrich *et al*., 2021; Yoast *et al*., 2021). Briefly, the cells were cultured on glass coverslips with 50-60% confluency. Next day the cells were washed with fresh media and incubated at 37°C for 45 min in culture media containing 4 μM Fura-2/acetoxymethyl ester (Molecular Probes, Eugene, OR, USA). Then the cells were washed and kept in HEPES-buffered saline solution (140 mM NaCl, 1.13 mM MgCl_2_, 4.7 mM KCl, 2 mM CaCl_2_, 10 mM D-glucose, and 10 mM HEPES, adjusted to pH 7.4 with NaOH) for at least 10 min before Ca^2+^ measurements were made. Time laps imaging was performed using Leica DMI8 microscope equipped with 40X oil objective. The change in fluorescence of Fura-2 was recorded and analyzed with Leica analysis software X. An ROI was drawn around each cell and 340/380 ratio of each cell was calculated.

### Endoplasmic reticulum Ca^2+^ measurements

Previously described method was used to measure endoplasmic reticular Ca^2+^ (ER Ca^2+^) (Emrich *et al*., 2021). Briefly, the cells were cultured at 50-60% confluency and transfected with 2-3 µg R-CEPIA_ER_. After 24-48 hrs, the cells were washed with HBSS (HEPES-buffered saline solution (140 mM NaCl, 1.13 mM MgCl_2_, 4.7 mM KCl, 2 mM CaCl_2_, 10 mM D-glucose, and 10 mM HEPES, adjusted to pH 7.4 with NaOH)) containing 2 mM CaCl_2_. The cells were stimulated with 300 µM ATP with no Ca^2+^. After complete release of ER Ca^2+^ the cells were stimulated with 2 mM Ca^2+^. The time laps images were acquired using Leica DMI8 microscope equipped with X 20 objective. The images were acquired every 40 sec intervals for 15-20 min. The fluorescent intensity was quantified using LAS X (Leica microsystems) software.

### Transmission Electron Microscopy (TEM)

As described previously (Pathak *et al*., 2020), briefly, the HCT116, STIM1KO, STIM2KO, and DKO HCT116 cells were cultured at 80-90% confluency. The cells were fixed with 1% glutaraldehyde in 0.1 M sodium phosphate buffer (pH 7.3). Then the fixed cells were washed with 100 mM Tris (pH 7.2) and 160 mM sucrose for 30 min. The cells were washed again twice with phosphate buffer (150 mM NaCl, 5 mM KCl, 10 mM Na_3_PO_4_, pH 7.3) for 30 min, followed by treatment with 1% OsO_4_ in 140 mM Na_3_PO_4_ (pH 7.3) for 1 hr. The cells were washed twice with water and stained with saturated uranyl acetate (C_4_H_6_O_6_U) for 1 hr, followed by dehydration in ethanol, and embedded in Epon (Electron Microscopy Sciences, Hatfield, PA). The embedded samples were cut in roughly 60 nm sections and stained with uranyl acetate and lead nitrate. The stained grids were imaged and analyzed using a Philips CM-12 electron microscope (FEI; Eindhoven, The Netherlands) and photographed with a Gatan Erlangshen ES1000W digital camera (Model 785, 4 k 3 2.7 k; Gatan, Pleasanton, CA). Mitochondrial morphology, and mitophagy was analyzed by double-blinded independent observers in at least 10 different micrographs per condition.

### Cell proliferation assays

As described previously (Pathak *et al*., 2020), briefly, the cells were harvested and 3000-5000 cells per well were plated in 96 well plate and kept at 37°C in 5% CO_2_ for 4 hr for proper adhesion. The CyQUANT-NF dye was diluted according to the manufacturer protocol and 100 µl/per well was added. After incubating the cells at 37 °C in 5% CO_2_ for 1 hr, fluorescence intensity (∼485 nm/∼530 nm) was measured using FlexStation 3 Multimode Plate Reader (VWR).

Normalized intensity= (I_t_-I_b_)\(I_0_-I_b_), I_t_= intensity at given time, I_b_= background intensity,I_0_= intensity at time zero

### Spheroid assays

Around 1000 cells were seeded in round bottom untreated 96 well plate with 200 µl media in each well (Nunclon Sphera 96U Well Round Bottom Plate). The plate was spin at 400 rpm for 2 min and kept at 37 °C, 5% CO_2_ incubator. Next day the cells were imaged under Leica DMI8 microscope equipped with 10X objective. It was considered as zero time point. Afterward the cells were imaged every 24 hr till the cells form large tight spheroid. The culture media of the spheroid was changed every 2 days. The images were analyzed with Image J.

### Colony formation assays

The cells were harvested, and 100 cells were cultured in each well of six well plates. The cells were incubated in 37 °C, 5% CO_2_ incubator for 6-7 days. The culture media was changed every two days. Finally, the colonies were stained with Colony fixation-staining solution, glutaraldehyde 6.0% (vol/vol), crystal violet 0.5% (wt/vol) in water. The colonies were imaged under FluorChem M gel imaging system. The number of colonies were counted using image J.

### In vitro migration and invasion assays

The migration and invasion were performed as described previously (Pathak *et al*., 2020). For the gap closure assay, the cell cycle was synchronized by starving the cells for 24 hr. Then 50,000 cells were cultured in ibidi-silicone insert with a defined cell-free gap. After 24 hr the inserts were lifted and the gap between two migrating fronts were measured at 0 hr, 12 hr, and 24 hr using Leica DMi8 equipped with 10X air objective. The area between the two migrating fronts was measured using ImageJ (NIH, Bethesda, USA).

% Gap closure= (A_0_-A_t_/A_0_)/100, A_0_= aria of gap measured immediately after lifting the insert, A_t_= aria of gap measured after indicated time of migration

For the transwell migration assay using, Briefly the cells were starved for 24 hr and 10,000 cells were plated on the FluoroBlok chamber in 200 µl media without FBS. The lower chamber was filled with 200 µl media with 10% FBS, which stimulated the cells to migrate towards the lower chamber. After 8 hr migration the upper face of the membrane was cleaned with cotton swab and lower part was stained with DAPI. The intensity of DAPI was measured using FlexStation 3 Multimode Plate Reader (VWR).

Invasion was measured using the BioCoat™ Tumor Invasion Plate with 8 µm pore size inserts. The upper face of the membrane was coated with growth factor reduced Matrigel. The cells were starved for 24 hr to synchronize the cell cycle and a total of 10,000 cells were plated on the chamber in 200 µl media without FBS. The cells were stimulated to invade towards preconditioned media with 10% FBS. After 24 hr invasion, the upper part of the membrane was cleaned with cotton swab and lower part was stained with DAPI and the intensity of DAPI was measured using FlexStation 3 Multimode Plate Reader (VWR).

%invasion= (Fluorescence of invading cells - Fluorescence of blank chamber)/ Fluorescence of migrating cells - Fluorescence of blank chamber) X100

### Extracellular acidification rate (ECAR) and Oxygen consumption rate (OCR)

Seahorse XFp Extracellular Flux Analyzer (Seahorse Bioscience) was used to measure ECAR, and OCR. Briefly, the cells were harvested, and 80,000-90,000 cells per well were cultured in a Seahorse XFp cell culture microplate with complete growth media and incubated in 37 °C, 5% CO_2_ incubator for 10-12 hr. The cells were washed with seahorse XF DMEM media (pH 7.4) and incubated in at 37 °C for 30 minutes. Further, the plate with cultured cells was placed in the Seahorse XFp Extracellular Flux Analyzer to record OCR and ECAR. For OCR measurements, oligomycin (1 µM), FCCP (p-trifluoromethoxy carbonyl cyanide phenylhydrazone-1 µM), and a mixture of Rotenone and antimycin A (Rote/AA-0.5 µM) were sequentially injected. Similarly, for the ECAR measurements, glucose (10 mM), oligomycin (1 µM), and 2-Deoxy Glucose (50 mM) were sequentially injected into each well at the indicated time points. The final values obtained for ECAR, and OCR, were normalized to protein amount. The data was analyzed by Seahorse XFp Wave software. The results for OCR are reported in pmols/minute/protein, and ECAR in mpH/minute/protein.

### Glucose and Lactate measurements

To measure glucose and lactate in media, briefly the cells were harvested, and 100,000 cells were plated in each well of 6 well plate with 2 ml of complete growth media. After 24 hr, 400 µl of the media was taken from one well of 6 well plate and divided into 4 wells of 96 well plate (quadruplicate for each STIM KO clone and respective control). Then the plate was kept in YSI 7100 multichannel biochemistry analyzer (YSI Life Sciences), to measure glucose and lactate levels in the media. Fresh media was used to measure basal level of glucose and lactate. After the measurement, the cells were harvested and number of cells was counted in each well of 6 well plate to make sure that the number of cells is same in each well. This experiment was performed independently at least thrice. Following formula was used to calculate glucose consumption and lactate generation-

Glucose consumption= (Glucose in fresh media-Glucose in the media with cells)/ total protein content

Lactate generation= (Lactate in the media with cells-Lactate in fresh media)/ total protein content

### Mitochondrial DNA Measurements

The mitochondrial DNS level was determined as described previously (Johnson *et al*., 2022). Briefly, the cells were washed, collected, and resuspended in DirectPCR Lysis Reagent (Viagen Biotech) and Proteinase K in 4:1 ratio. Total DNA was isolated using isopropanol and precipitated with 70% ethanol precipitation method and washed three times. A RT-qPCR (50 °C 2-min activation step, a 95 °C 2-min melt step, and 40 cycles of 95°C for 15 s followed by 56 °C for 15 s and 72 °C for 30 s), was performed. Three sets of specific primers for mitochondrial DNA and two sets for genomic DNA were used. The comparative Ct method was used to quantify mitochondrial DNA copy number.

### Mitochondrial burden, Mitochondrial ROS, Cellular ROS, and Mitochondrial membrane potential measurements

To determine mitochondrial burden in the CRC cells, 1,000,000 cells were stained with 100 nM Mito tracker green. The cells were incubated in the complete growth media and kept at 37°C in 5% CO2 for 15 min, and the intensity of staining was measured using LSRII flow cytometer using FACSDiva software (BD Biosciences) and analyzed with FlowJo software (Tree Star).

As previously described (Pathak *et al*., 2020) the cells were cultured in 6-well plates at 50–60% confluency. To measure cellular ROS and mitochondrial ROS, 1,000,000 cells were stained with CellRox (according to manufacturer protocol) and MitoSOX (2.5 µM in HBSS at 37°C for 45 min in absence of CO_2_) respectively. Antimycin (50 µM) was used as a positive control. The cells were washed with warm HBSS before the intensity measurement.

For mitochondrial membrane potential measurement, 1,000,000 cells were harvested and loaded with Tetramethyl rhodamine (TMRM) dye. The cells were stained with 100 nM TMRM dye in complete growth media and kept at 37°C in 5% CO2 for 30 min. Carbonyl cyanide m-chlorophenyl hydrazone (CCCP-50 µM) was used as a positive control. The cells were washed with warm complete growth media before intensity measurement. The intensity of staining was measured using LSRII flow cytometer using FACSDiva software (BD Biosciences) and analyzed with FlowJo software (Tree Star).

### Quantification and statistical analysis

The data are represented as mean ±SEM and analyzed using Origin pro 2019b 64 bit (Origin lab). In the box plot, the box represents 25^th^ to 75^th^ interquartile range, midline in the box represents median and solid square box represents mean data points. To test single variables between two groups Student’s t test, paired t-test or Kruskal-Wallis ANOVA was performed (mentioned in the figure legend). One-way ANOVA followed by post hoc Tukey’s test was used for comparison between multiple groups versus the control group. A paired t-test was used to compare between two groups. The p value < 0.05 was considered to be significant and is presented as * p < 0.05, ** p < 0.01, or *** p < 0.001.

### Contact for Reagent and Resource sharing

All the request and queries about reagents and resources should be directed to and fulfilled by lead contact Dr. Mohamed Trebak.

## Acknowledgments

The authors acknowledge Dr. Han Chen from The Pennsylvania State University College of Medicine EM facility for assistance with TEM imaging and Dr. Katherine Aird for generously sharing the YSI analyzer. This study was supported by the National Heart, Lung, and Blood Institute (R01-HL123364, R01-HL097111, and R35-HL150778 to MT), National Institute on Aging (R21-AG050072 to MT), and National Heart, Lung, and Blood Institute (K99 HL163403-01 to TP). GSY and WAK are supported by the Peter and Marshia Carlino Fund for IBD research. We also acknowledge the Flow Cytometry and Cell Sorting, Imaging, Informatics and Data Analysis Core facility (The Pennsylvania State University, College of Medicine and The University of Pittsburgh).

## Author contributions

T.P.- Conceptualization, Data curation, Formal analysis, Investigation, Visualization, Methodology, Writing - original draft, Writing - review and editing

J.C.B.- Data curation, Formal analysis, Writing - review and editing

M.T.J.- Data curation, Formal analysis

P.X.- Data curation, Formal analysis,

V.W.- Resources, Software, Formal analysis

W.K.- Resources

G.Y.- Resources, Writing - review and editing

N.H.- Resources, Writing - review and editing

M.T.- Conceptualization, Supervision, Funding acquisition, Methodology, Project administration, Writing - review and editing

## Legends to supplementary figures

**Supplementary Figure 1.**
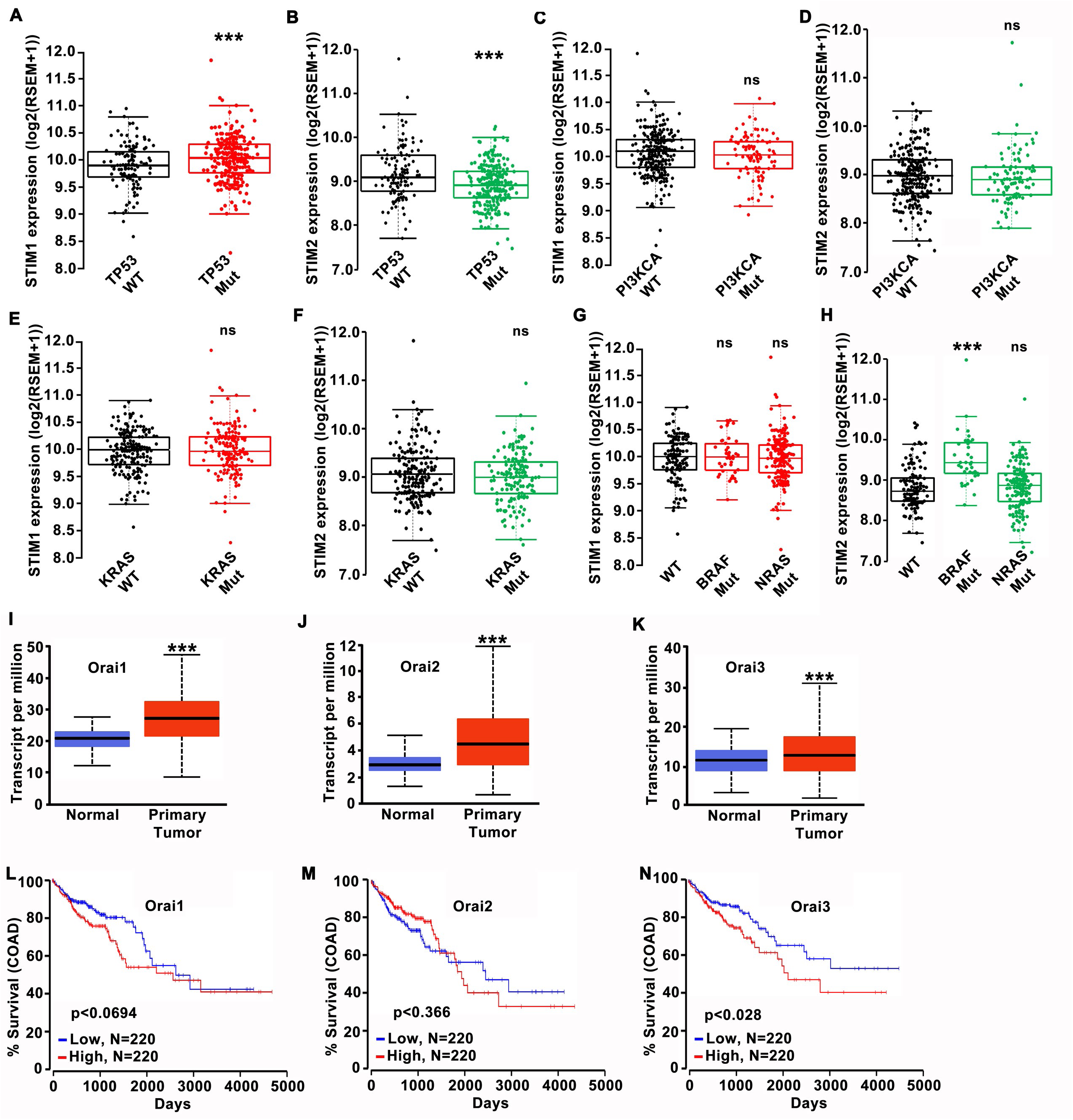
Loss of STIM2 is associated with poor prognosis of CRC patients. (A, B) TCGA data showing (A) STIM1 and (B) STIM2 mRNA levels in normal and TP53 mutated conditions. (C, D) TCGA data showing (C) STIM1 and (D) STIM2 mRNA levels in normal and PI3KCA mutated conditions. (E, F) TCGA data showing (E) STIM1 and (F) STIM2 mRNA levels in normal and KRAS mutated conditions. (G, H) TCGA data showing (G) STIM1 and (H) STIM2 mRNA levels in normal and BRAF and NRAS mutated conditions. (I-K) TCGA data showing (I) Orai1, (J) Orai2, and (K) Orai3 mRNA levels in the tumor and adjacent normal tissue. (L-N) Kaplan plot for (L) Orai1, (M) Orai2, and (N) Orai3 showing survival of colon adenocarcinoma (COAD) patient. Kruskal-Wallis ANOVA was performed to test single variables between the two groups. *p<0.05, **p<0.01, and ***p<0.001.

**Supplementary Figure 2.**
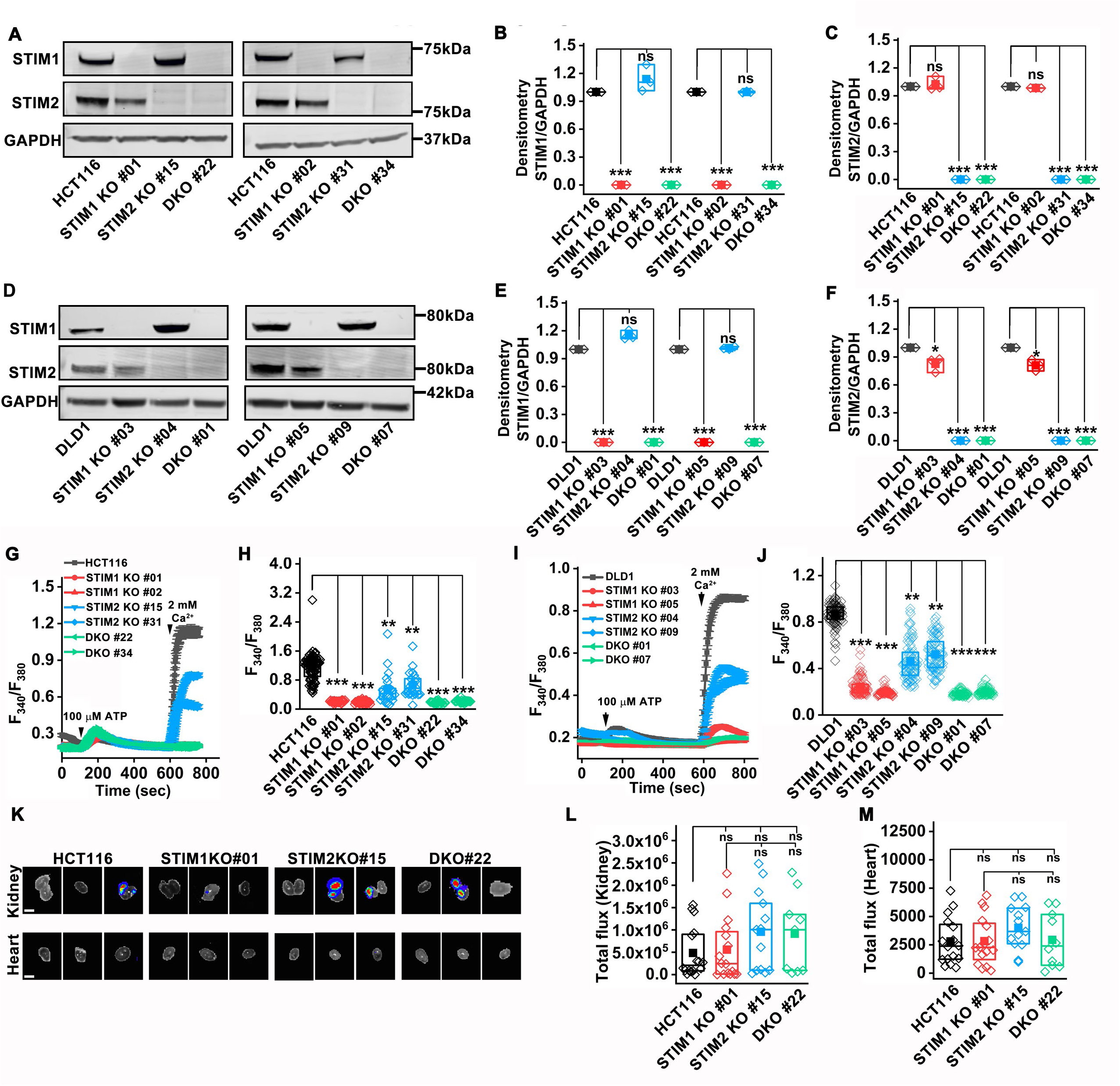
Loss of STIM2 leads to a partial reduction in SOCE in CRC cells. (A-C) Measurement of STIM1 and STIM2 protein level (A) representative western blot probed with anti-STIM1, STIM2, and GAPDH antibody, and densitometric analysis of (B) STIM1, and (C) STIM2 normalized to GAPDH in HCT116 and STIM1 KO, STIM2 KO, and DKO clones of HCT116 cells. (D-F) Measurement of STIM1 and STIM2 protein level (D) representative western blot probed with anti-STIM1, STIM2, and GAPDH antibody, and densitometric analysis of (E) STIM1, and (F) STIM2 normalized to GAPDH in DLD1 and STIM1 KO, STIM2 KO, and DKO clones of DLD1 cells. (G) Cytosolic Ca^2+^ measurements with Fura-2 in response to 100 µM ATP applied first in nominally Ca^2+^-free external solution and subsequently with 2 mM external Ca^2+^. The traces are represented as mean ± S.E.M (H) Quantification of peak SOCE after addition of 2 mM Ca^2+^ in HCT116 (n = 82) and STIM1 KO #01 (n=34), STIM1 KO #02 (n=43), STIM2 KO #15 (n=23), STIM2 KO #31 (n=24), DKO #22 (n=20) and DKO #34 (n=30) clones of HCT116 cells. (I) Fura-2 Ca^2+^ measurements upon store depletion with 100 µM ATP in 0 mM Ca^2+^ and subsequent addition of 2 mM Ca^2+^. (J) Quantification of peak SOCE after addition of 2 mM Ca^2+^ in DLD1(n = 90) and STIM1 KO #03 (n=90), STIM1 KO #05 (n=90), STIM2 KO #04 (n=90), STIM2 KO #09 (n=90), DKO #01 (n=90) and DKO #07 (n=90) clones of DLD1 cells. (K-M) Post intrasplenic injection of HCT116, STIM1 KO #01, STIM2 KO #15, and DKO #22 cells into the NOD SCID mice, the organs were harvested, and luminance was measured (K) representative image of kidney and heart, and quantification of total luminance in (L) kidney, and (M) heart. (Scale bar 0.5 cm) All experiments were performed ≥three times with similar results. Statistical significance was calculated using paired t-test. *p<0.05, **p<0.01, ***p<0.001

**Supplementary Figure 3.**
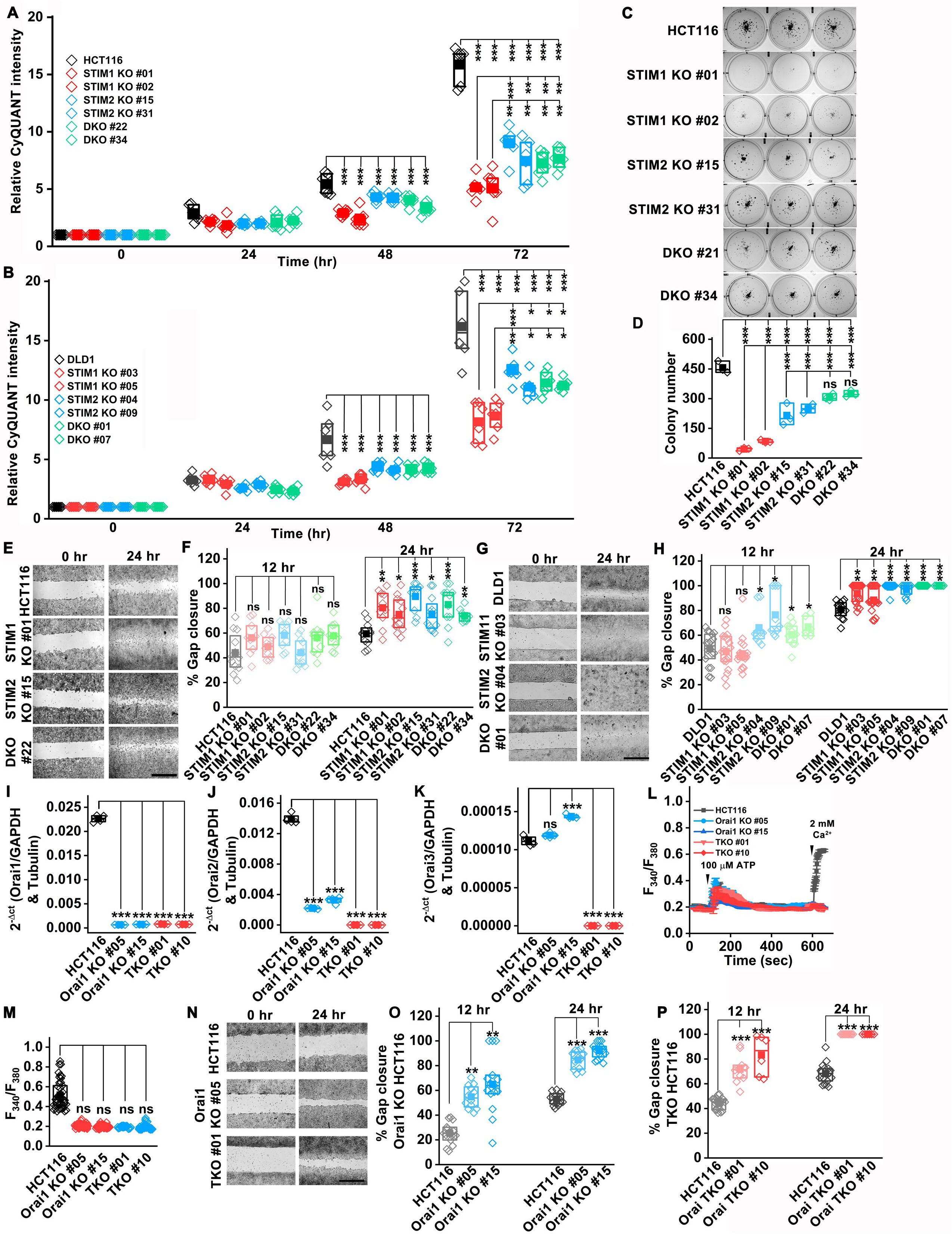
Loss of STIM2 promotes migration of CRC cells. (A, B) The proliferation assay was performed by staining (A) HCT116 and STIM1 KO, STIM2 KO, and DKO clones of HCT116 cells, and (B) DLD1 and STIM1 KO, STIM2 KO, and DKO clones of DLD1 cells using CyQUANT dye. (C, D) Representative bright field image of (C) colony formation assay and (D) quantification of the number of colonies formed by HCT116 and STIM1 KO, STIM2 KO, and DKO clones of HCT116 cells. (E, F) In vitro migration assay performed using HCT116 and STIM1 KO, STIM2 KO, and DKO clones of HCT116 cells. (E) The image showing gap closure at 0 and 24 hr post culture, and (F) quantification of percent gap closure. Scale bar 1 mm. (G, H) In vitro migration assay performed using DLD1 and STIM1 KO, STIM2 KO, and DKO clones of DLD1 cells. (G) The image showing gap closure at 0 and 24 hr post culture, and (H) quantification of percent gap closure. Scale bar 1 mm. (I-K) RT-qPCR analysis of mRNA levels of (I) Orai1, (J) Orai2, and (K) Orai3 in HCT116 and Orai1 KO and Orai TKO clones of HCT116. (L, M) Fura-2 Ca^2+^ measurements upon store depletion with 100 µM ATP in 0 mM Ca^2+^ and subsequent addition of 2 mM Ca^2+^. (L) The trace represents mean± S.E.M. (M) Quantification of peak SOCE after addition of 2 mM Ca^2+^ in HCT116 (n = 41) and Orai1 KO #05 (n=58), Orai1 KO #15 (n=68), TKO #01 (n=30), and TKO #10 (n=71) clones of HCT116 cells. (N) Representative image of in vitro migration assay performed using HCT116 and Orai1 KO, and Orai TKO clones of HCT116 cells. (Scale bar 1 mm) (O, P) The quantification of percent gap closure by (O) HCT116 and Orai1 clones of HCT116 and (P) HCT116 and TKO clones of HCT116. All experiments were performed ≥three times with similar results. Statistical significance was calculated using paired t-Test was used to compare between groups. *p<0.05, **p<0.01, ***p<0.001

**Supplementary Figure 4.**
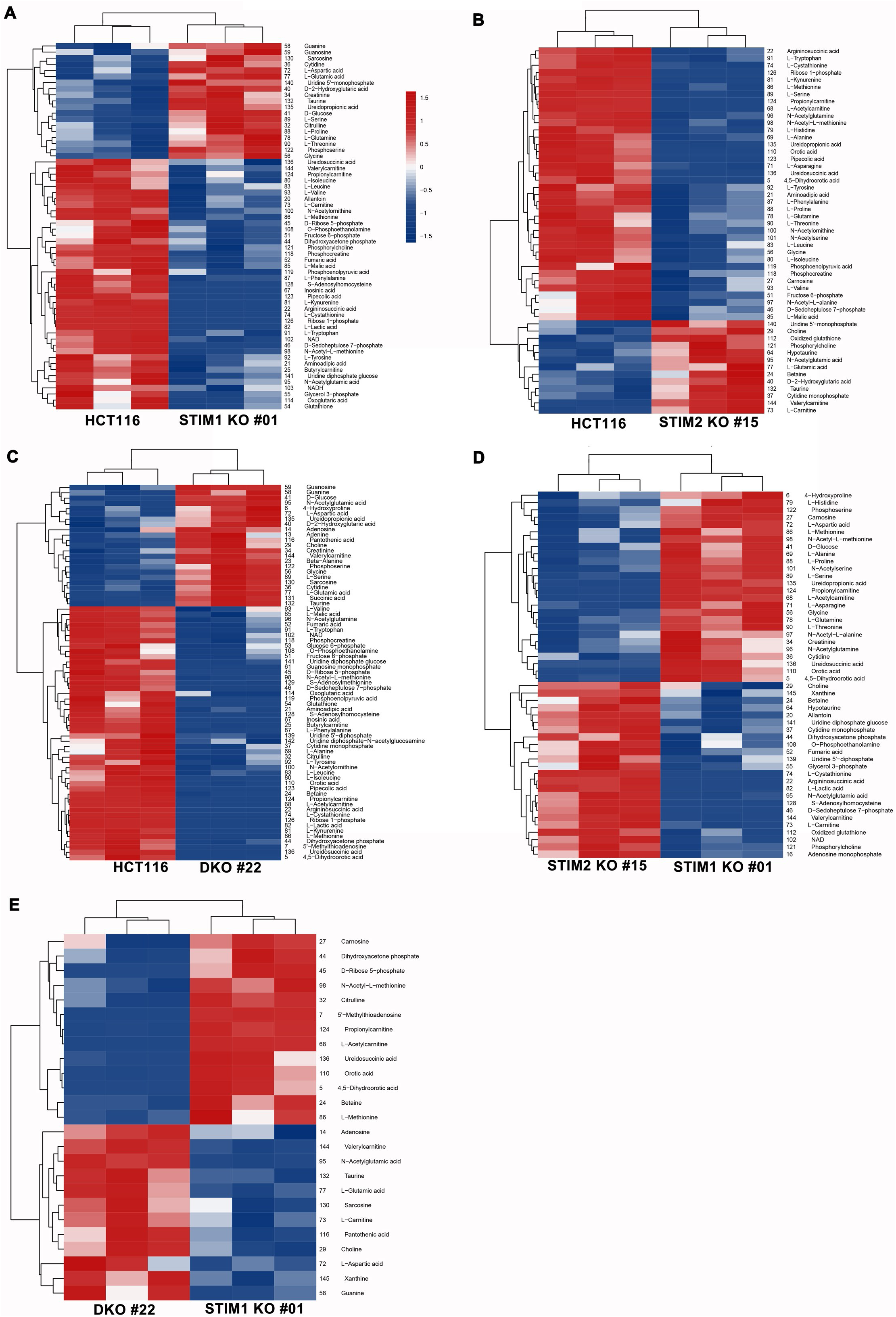
Loss of STIM2 causes metabolic rewiring of CRC cells. (A-E) The levels of polar metabolites were measured using whole-cell metabolomics. Heatmap represents comparison of polar metabolites levels between (A) HCT116 and STIM1 KO #01 clone, (B) HCT116 and STIM2 KO #15 clone (C) HCT116 and DKO #22 clone, (D) STIM2 KO #15 and STIM1 KO #01 clone of HCT116, and (E) DKO #22 and STIM1 KO #01 clone of HCT116.

**Supplementary Figure 5.**
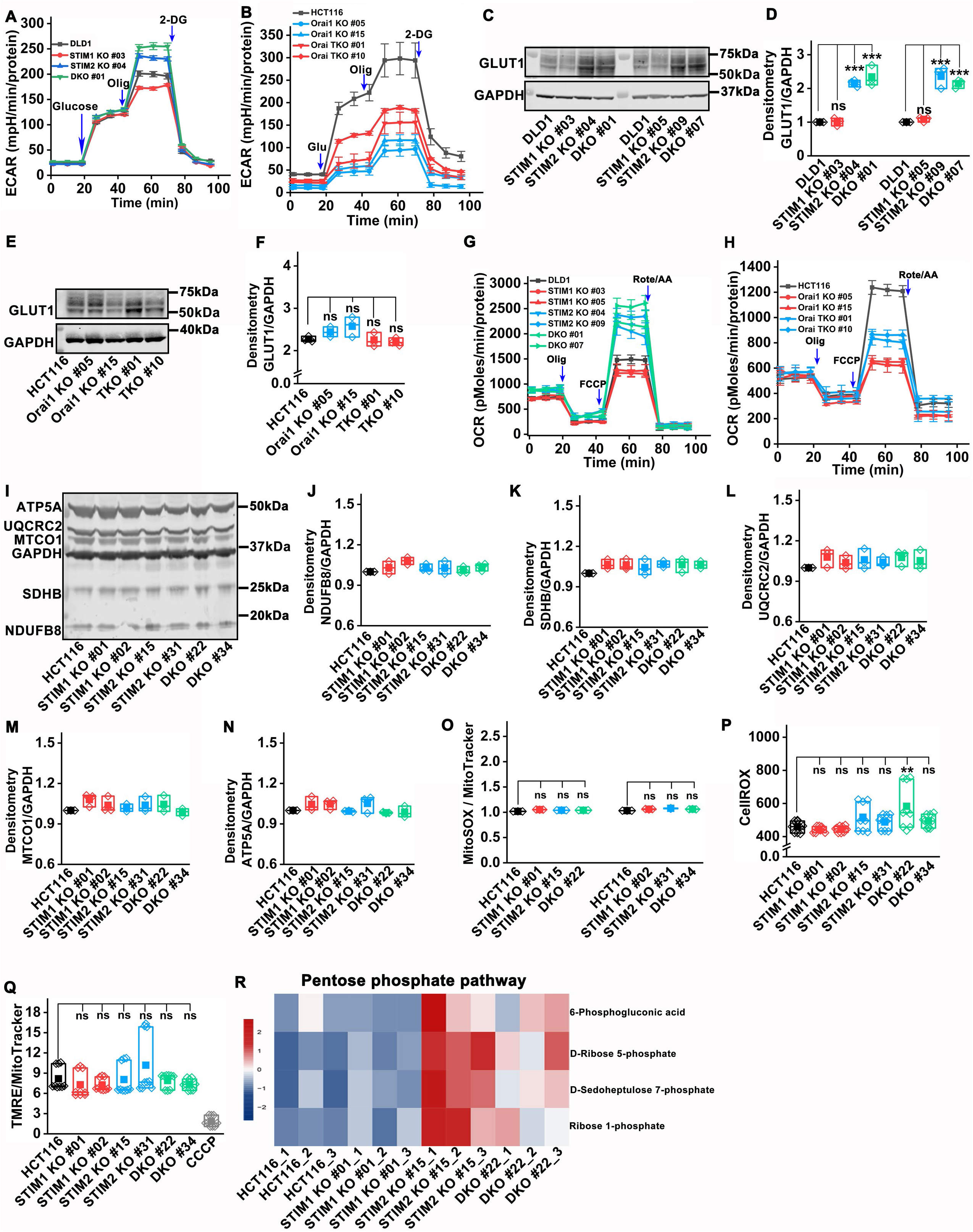
STIM2 regulates metabolism of CRC cells. (A) Measurement of ECAR in DLD1, STIM1KO #03, STIM2KO #04, and DKO #01 clones of HCT116. (B) ECAR measurement in HCT116, and Orai1 KO and TKO clones of HCT116. (C, D) GLUT1 protein level measurement using western blot. (C) Representative western blot probed with anti-GLUT1 and GAPDH antibody, and (D) densitometric quantification of GLUT1 levels in DLD1, STIM1 KO, STIM2 KO, and DKO clones of DLD1. (E, F) The measurement of GLUT1 protein levels using (E) western blot and (F) densitometric analysis in HCT116 and Orai1 KO and TKO clones of HCT116. (G, H) OCR measurement in (G) DLD1 and STIM1 KO, STIM2 KO and DKO, clones of DLD1cells, and (H) HCT116 and Orai1 KO, and TKO clones of HCT116. (I) Quantification of components of electron chain complex components. The representative western blot probed with a cocktail of antibodies against NADH dehydrogenase [ubiquinone] 1 b subcomplex subunit 8 (NDUFB8, Complex-I), Succinate dehydrogenase [ubiquinone] iron-sulfur subunit (SDHB, Complex-II), Cytochrome b-c1 complex subunit 2 (UQCRC2, Complex-III), Cytochrome c oxidase subunit 1 (MTCO1, Complex IV), and ATP synthase subunit an (ATP5A, Complex-V). (J-N) Densitometric analysis of protein levels of (J) NDUFB8, (K) SDHB, (L) UQCRC2, (M) MTCO1, and (N) ATP5A in HCT116 and STIM1 KO, STIM2 KO, and DKO clones of HCT116. (O) Measurement of mitochondrial ROS in HCT116 and STIM1 KO, STIM2 KO, and DKO clones of HCT116 using flow cytometry. The cells were stained with MitoSOX and MitoTracker. The graph represents MitoSOX intensity normalized to MitoTracker. (P) Cytosolic ROS measurement in HCT116 and STIM1 KO, STIM2 KO, and DKO clones of HCT116 using flow cytometry. The cells were stained with CellROX and the graph represents CellROX intensity. (Q) Mitochondrial membrane potential measurement in HCT116 and STIM1 KO, STIM2 KO, and DKO clones of HCT116 using flow cytometry. The cells were stained with TMRE and MitoTracker. The graph represents TMRE intensity normalized to MitoTracker. (R) Heatmap showing levels of pentose phosphate pathway metabolites in HCT116 and STIM1 KO, STIM2 KO, and DKO clones of HCT116. All the experiments were performed ≥three times with similar results. The statistical significance was calculated using paired t-Test. *p<0.05, **p<0.01, ***p<0.001

**Supplementary Figure 6.**
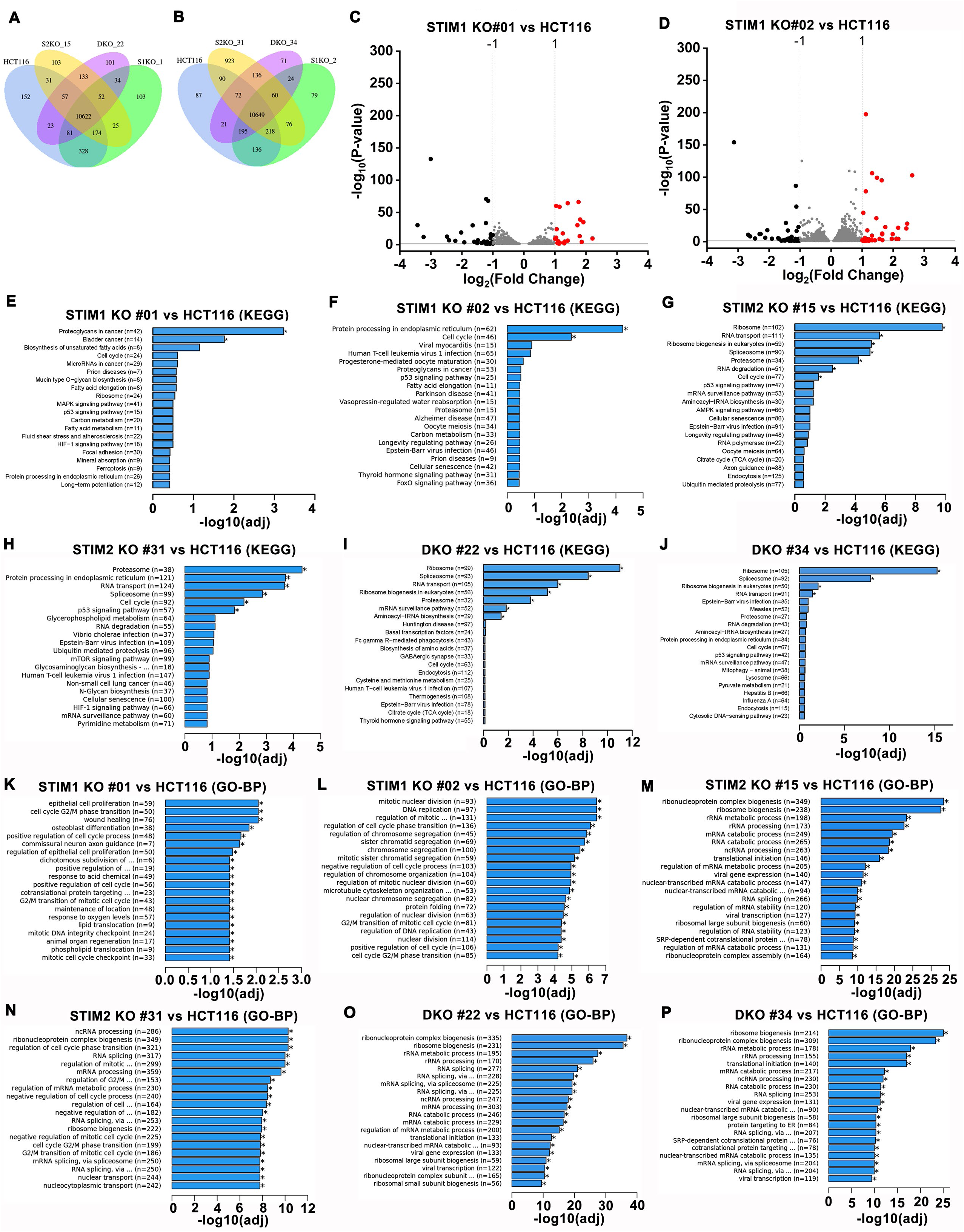
Loss of STIM2 induces ribosome biogenesis. (A, B) Venn diagram showing common and uniquely expressed genes between (A) HCT116, and STIM1 KO #01, STIM2 KO#15, DKO #22 clones of HCT116, and (B) HCT116 and STIM1 KO #02, STIM2 KO #31, and DKO #34 clones of HCT116. (C, D) Volcano plot showing differentially expressed genes between (C) HCT116 and STIM1 KO #01 clone of HCT116 and (D) HCT116 and STIM1 KO #02 clone of HCT116. (E-J) The Kyoto Encyclopedia of Genes and Genomes (KEGG) analysis showing enrichment of pathways based on molecular interactions. The plot shows the top 20 pathways that are significantly enriched in (E) STIM1 KO #01, (F) STIM1 KO #02, (G) STIM2 KO #15, (H) STIM2 KO #31, (I) DKO #22, and (J) DKO #34 clones of HCT116 compared to wildtype HCT116 cells. (K-P) The Gene ontology (GO) analysis showing enrichment based on biological process (BP). The plot showing the top 20 pathways that are significantly enriched in (K) STIM1 KO #01, (L) STIM1 KO #02, (M) STIM2 KO #15, (N) STIM2 KO #31, (O) DKO #22, and (P) DKO #34 clones of HCT116 compared to wildtype HCT116 cells.

**Supplementary Figure 7.**
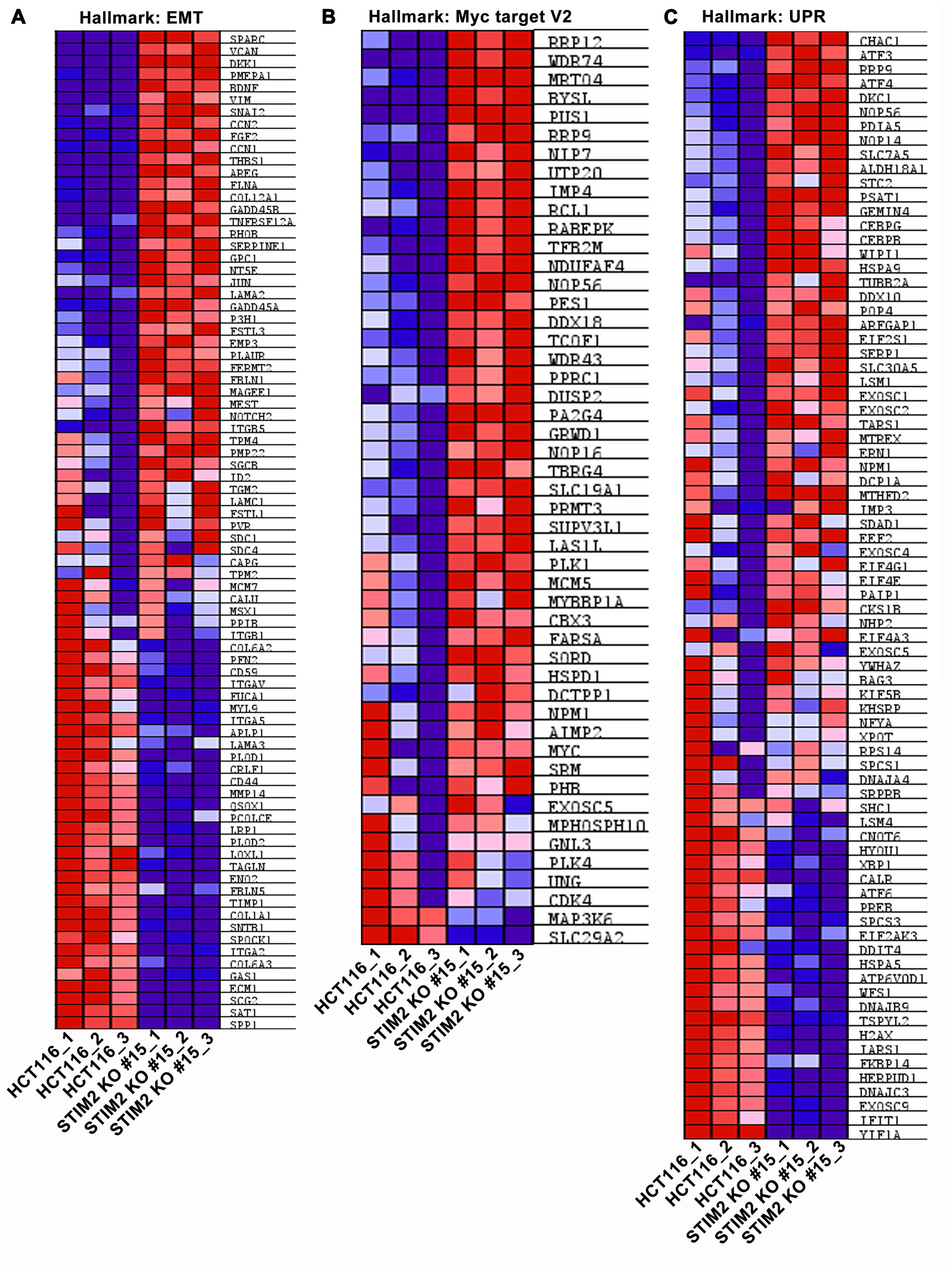
Loss of STIM2 induces expression of EMT and UPR hallmark genes. (A, C) Heatmap showing differential expression of genes of hallmark (A) EMT, (B) Myc target version 2, and (C) unfolded protein response (UPR) between HCT116 and STIM2 KO #15 clones of HCT116 cells.

**Supplementary Figure 8.**
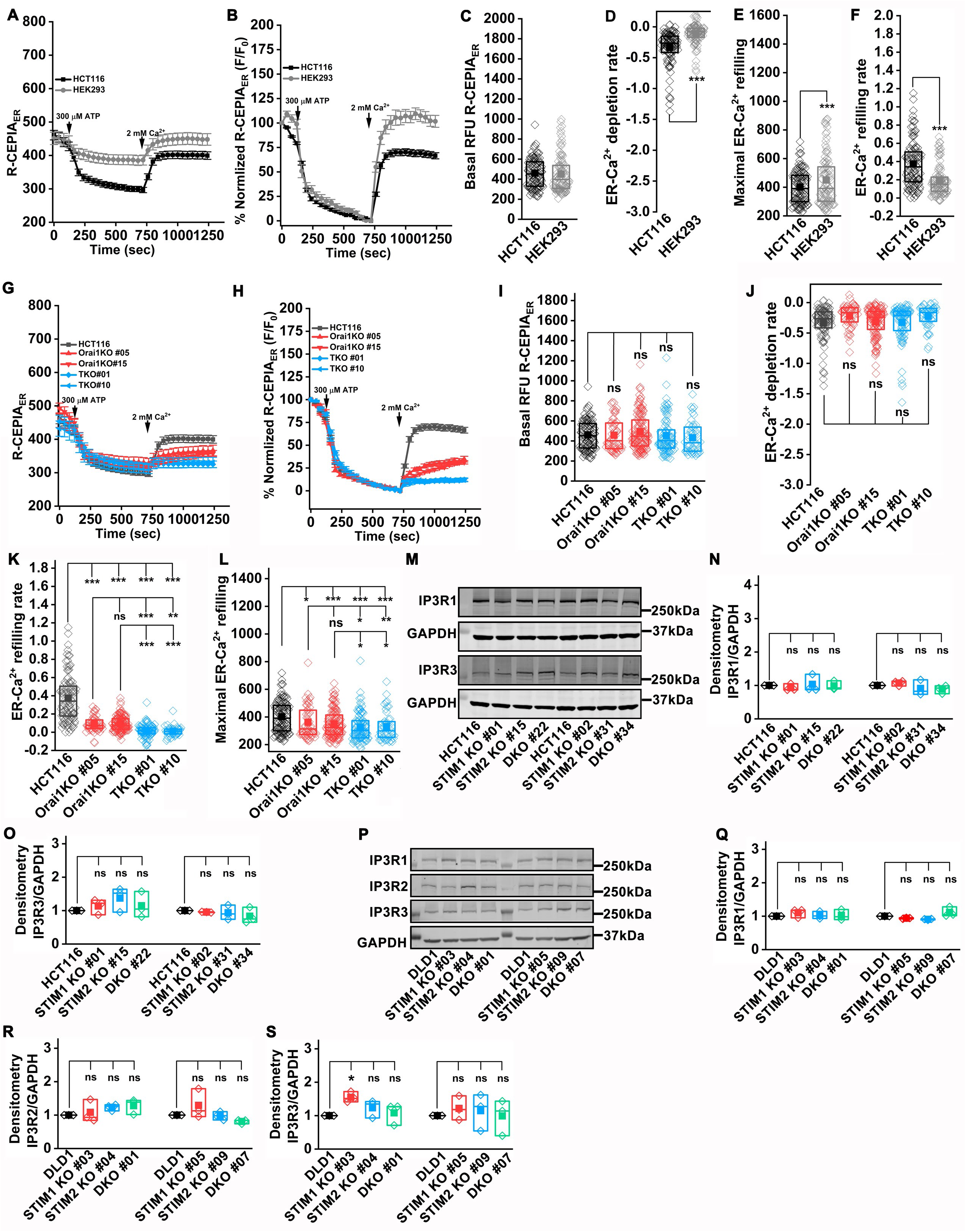
Loss of Orai does not affect ER Ca^2+^ levels in HCT116 cells. (A, B) ER Ca^2+^ measurement using genetically encoded R-CEPIA_ER_. The ER Ca^2+^ depletion was stimulated with 300 µM ATP in 0 mM Ca^2+^ for 10 minutes, followed by the addition of 2 mM Ca^2+^. (A) The graph represents the mean ± S.E.M. of R-CEPIA_ER_ fluorescence in HCT116 and, HEK293 cells and (B) Percentage of normalized F/F_0_ value of R-CEPIA_ER_ in HCT116 and HEK293 cells. The traces are represented as mean ± S.E.M (C-F) Quantification of (C) Basal ER Ca^2+^, (D) ER Ca^2+^ depletion rate 3 min post stimulation in 0 Ca^2+^, (E) maximal ER Ca^2+^ at 15 min in 2 mM Ca^2+^, and (F) ER Ca^2+^ refilling rate at 12.5 min in 2 mM Ca^2+^ in HCT116 (n=150), and HEK293 (n=152). (G, H) ER Ca^2+^ measurement using genetically encoded R-CEPIA_ER_. The ER Ca^2+^ depletion was stimulated with 300 µM ATP in 0 mM Ca^2+^ for 10 minutes, followed by the addition of 2 mM Ca^2+^. The graph represents the mean ± S.E.M. of (G) HCT116, Orai1 KO #05, Orai1 KO #15, TKO #01, and TKO #10 clone of HCT116, and (H) Percentage of normalized F/F_0_ value of R-CEPIA_ER_ in HCT116, Orai1 KO #05, Orai1 KO #15, TKO #01, and TKO #10 clone of HCT116 cells. The traces are represented as mean ± S.E.M (I-L) Quantification of (I) Basal ER Ca^2+^, (J) ER Ca^2+^ depletion rate 3 min post stimulation in 0 Ca^2+^, (K) ER Ca^2+^ refilling rate at 12.5 min in 2 mM Ca^2+^, and (L) maximal ER Ca^2+^ at 15 min in 2 mM Ca^2+^ in in HCT116 (n=150), Orai1 KO #05 (n=124), Orai1 KO #15 (n=95), TKO #01 (n=80), TKO #10 (n=93). (M-O) Measurement of IP3R1 and IP3R3 protein level by probing the western blots with anti-IP3R1, anti-IP3R3 and GAPDH antibody (M) representative blot, (N) densitometric analysis of IP3R1 normalized to GAPDH and (O) densitometric analysis of IP3R3 normalized to GAPDH in HCT116 and clones of STIM1 KO, STIM2 KO, and DKO in HCT116 cells. (P-S) Measurement of IP3R1, IP3R2 and IP3R3 protein level by probing the western blots with anti-IP3R1, anti-IP3R2, anti-IP3R3 and GAPDH antibody (P) representative blot, (Q) densitometric analysis of IP3R1, (R) IP3R2, and (S) IP3R3 normalized to GAPDH in DLD1 and clones of STIM1 KO, STIM2 KO, and DKO in DLD1 cells. ER Ca^2+^ measurements from HCT116, HEK293, Orai1 KO, and Orai TKO were performed together. All the experiments were performed ≥three times with similar results. The statistical significance was calculated using paired t-test. *p<0.05, **p<0.01, ***p<0.001

**Supplementary figure 9.**
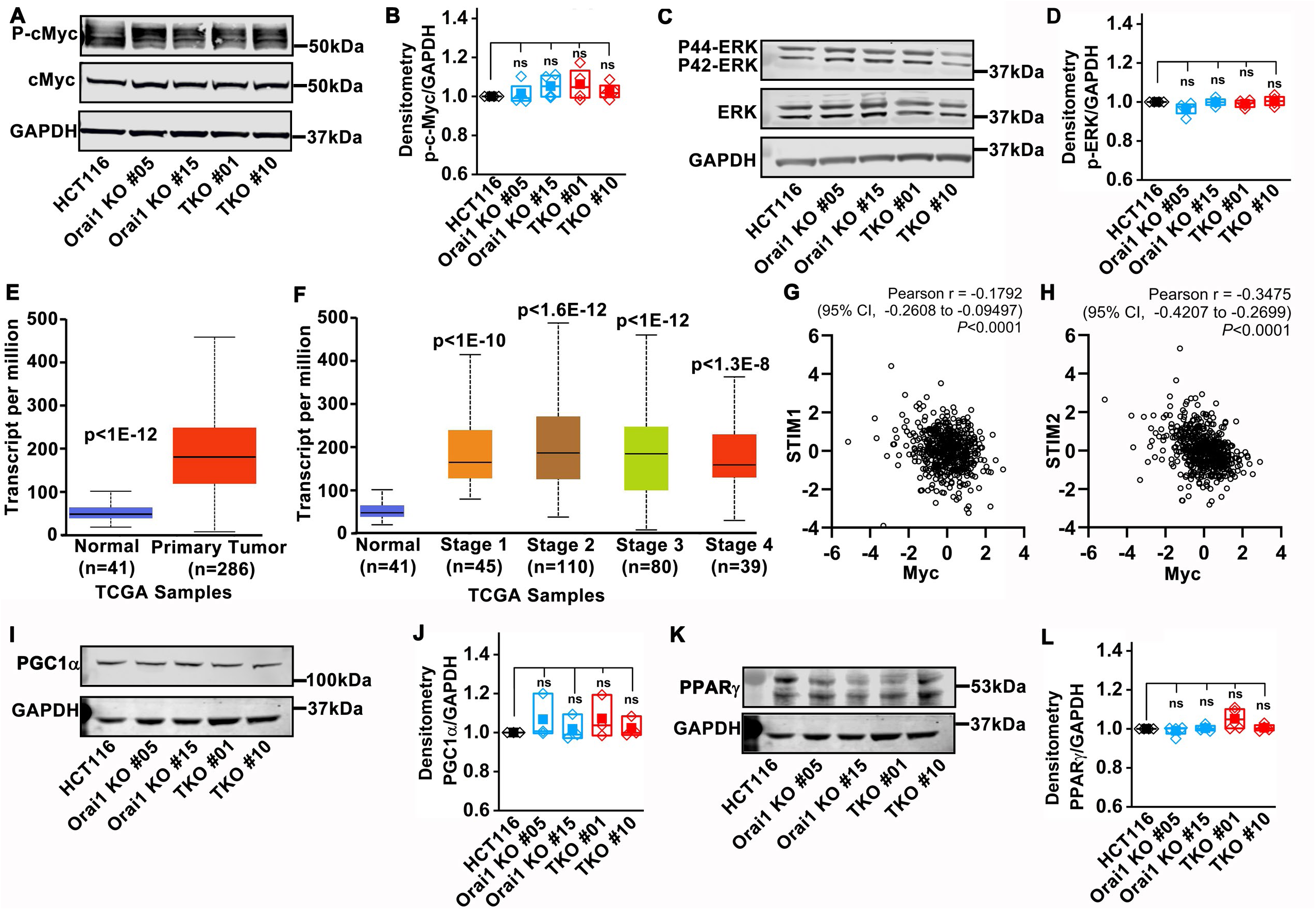
Deletion of Orai did not affect cMyc, ERK, and PPARɣ levels. (A, B) Quantification cMyc levels using (A) western blot probed with anti-phospho-cMyc, cMyc, and GAPDH and (B) densitometric analysis of phospho-cMyc in HCT116, Orai1 KO, and TKO clones of HCT116. (C, D) Quantification of phospho-ERK levels using (C) western blot probed with anti-phospho-ERK, ERK, and GAPDH and (D) densitometric analysis of phospho-ERK in HCT116, Orai1 KO, and TKO clones of HCT116. (E-F) TCGA analysis using ULCAN showing cMyc mRNA levels in (E) normal and primary tumors, (F) normal and individual stages of CRC growth. (G, H) Correlation of cMyc mRNA expression with STIM1 and STIM2 mRNA expression in Colorectal Adenocarcinoma samples from TCGA (n=524, Pan Cancer Atlas). Data obtained using cBioPortal and expressed as z-scores relative to all samples (log RNA Seq V2 RSEM). (I-L) PGC1α and PPARɣ levels quantified using western blot probed with (I) anti-PGC1α, (K) anti-PPARɣ and GAPDH antibody, and densitometric analysis of (J) PGC1α, and (L) PPARɣ levels in HCT116, Orai1 KO, and TKO clones of HCT116. All experiments were performed ≥three times with similar results. Statistical significance was calculated using paired t-test. *p<0.05, **p<0.01, ***p<0.001

**Supplementary figure 10.**
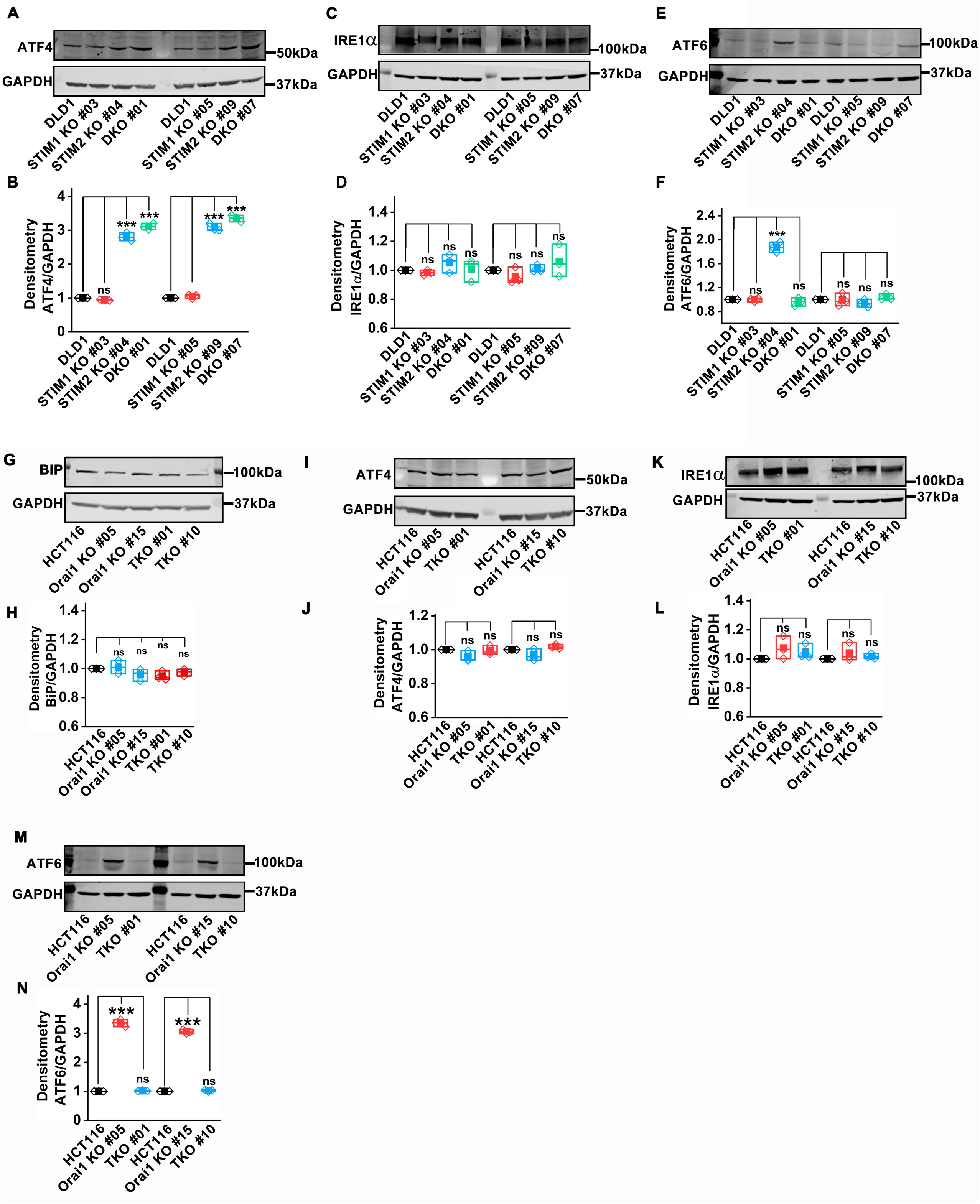
Loss of Orai did not activate the ER stress response in CRC cells. (A, B) Quantification of ATF4 levels using (A) western blot probed with anti-ATF4 and GAPDH antibody, and (B) densitometric analysis in DLD1, STIM1 KO, STIM2 KO, and DKO clones of DLD1. (C, D) Quantification of IRE1α levels using (C) western blot probed with anti-IRE1α and GAPDH antibody, and (D) densitometric analysis in DLD1, STIM1 KO, STIM2 KO, and DKO clones of DLD1. (E, F) Quantification of ATF6 levels using (E) western blot probed with anti-ATF6 and GAPDH antibody, and (F) densitometric analysis in DLD1, STIM1 KO, STIM2 KO, and DKO clones of DLD1. (G, H) Quantification of BiP levels using (G) western blot probed with anti-Bip and GAPDH antibody, and (H) densitometric analysis in HCT116, Orai1 KO, and TKO clones of HCT116. (I, J) Quantification of ATF4 levels using (I) western blot probed with anti-ATF4 and GAPDH antibody, and (J) densitometric analysis in HCT116, Orai1 KO, and TKO clones of HCT116. (K, L) Quantification of IRE1α levels using (K) western blot probed with anti-IRE1α and GAPDH antibody, and (L) densitometric analysis in HCT116, Orai1 KO, and TKO clones of HCT116. (M, N) Quantification of ATF6 levels using (M) western blot probed with anti-ATF6 and GAPDH antibody, and (N) densitometric analysis in HCT116, Orai1 KO, and TKO clones of HCT116. All experiments were performed ≥three times with similar results. Statistical significance was calculated using paired t-test. *p<0.05, **p<0.01, ***p<0.001

